# Two circadian oscillators in one cyanobacterium

**DOI:** 10.1101/2021.07.20.453058

**Authors:** Christin Köbler, Nicolas M. Schmelling, Alice Pawlowski, Philipp Spät, Nina M. Scheurer, Kim Sebastian, Lutz C. Berwanger, Boris Maček, Anika Wiegard, Ilka M. Axmann, Annegret Wilde

**Author notes:** ^1^C.K., N.M.S, and A.P. contributed equally to this study. Corresponding authors: Prof. Dr. Annegret Wilde, Albert-Ludwigs-Universität Freiburg, Institut für Biologie III, Schänzlestr. 1, 79104 Freiburg, Germany, *Prof. Dr. Ilka Axmann, Institute for Synthetic Microbiology, Biology Department, Heinrich Heine University Düsseldorf, 40225 Düsseldorf, Germany, **Email:**. **Author Contributions:** C.K., I.M.A. and A.W. designed the study. C.K., N.M.S., A.P., P.S., N.M.Sche., K.S., A. Wie, and L.B. performed and analyzed the experiments. All authors interpreted and discussed the data. C.K., N.M.S., A.P., P.S., B.M., A. Wie, I. M.A., and A.W. wrote the paper.

## Abstract

Organisms from all kingdoms of life have evolved diverse mechanisms to address the predictable environmental changes resulting from the Earth’s rotation. The circadian clock of cyanobacteria is a particularly simple and elegant example of a biological timing mechanism for predicting daily changes in the light environment. The three proteins KaiA, KaiB, and KaiC constitute the central timing mechanism that drives circadian oscillations in the cyanobacterium *Synechococcus elongatus* PCC 7942. In addition to the standard oscillator, *Synechocystis* sp. PCC 6803, another model organism for cyanobacterial research, harbors several divergent clock homologs. Here, we describe a potential new chimeric KaiA homolog that we named KaiA3. At the N-terminus, KaiA3 is similar to the NarL-type response regulator receiver domain. However, its similarity to canonical NarL transcription factors drastically decreases in the C-terminal domain, which resembles the circadian clock protein, KaiA. In line with this, we detected KaiA3-mediated stimulation of KaiC3 phosphorylation. Phosphorylation of KaiC3 was rhythmic over 48 h in vitro in the presence of KaiA3 and KaiB3 as well as in *Synechocystis* cells under free-running conditions after light/dark entrainment. This results in the presence of two different oscillators in a single-celled prokaryotic organism. Deletion of the *kaiA3* gene leads to KaiC3 dephosphorylation and results in growth defects during mixotrophic growth and in the dark. In summary, we suggest that KaiA3 is a nonstandard KaiA homolog, thereby extending the KaiB3-KaiC3 system in Cyanobacteria and potentially other prokaryotes.

## Introduction

The three genes, *kaiA*, *kaiB,* and *kaiC,* encode the core circadian oscillator in Cyanobacteria^1^. Over the last few decades, the biochemical interplay between these three proteins has been studied in great detail in *Synechococcus elongatus* PCC 7942 (hereafter *Synechococcus*). The KaiC protein forms a homohexamer and has autokinase, autophosphatase, and ATPase activities^1–4^. By associating with KaiC, KaiA stimulates the autokinase and ATPase activities of KaiC, and thus, the protein gets phosphorylated^5–7^. Upon phosphorylation of two neighboring residues (Ser431 and Thr432), KaiC undergoes structural rearrangements, exposing a binding site for KaiB^8–10^. After binding, KaiB sequesters KaiA from KaiC, promoting KaiC’s autophosphatase activity, and the protein reverts back to its unphosphorylated state^8, 9, 11^. The interplay between KaiA and KaiB is crucial for the KaiC phosphorylation cycle, which confers clock phase and rhythmicity to the cell^12, 13^. For a more detailed review on the KaiABC oscillator and its regulatory network, see Cohen and Golden^14^, Swan *et al*.^15^ and Snijder and Axmann^16^.

Although most studies on prokaryotic circadian rhythms have focused on the cyanobacterium *Synechococcus*, it has been shown that the standard KaiABC system is functionally conserved in other cyanobacteria^17^. However, in addition to the standard KaiABC system, divergent homologs of KaiB and KaiC have been identified in cyanobacteria, other bacterial species, and archaea^18^. The structure, mechanism of function, and physiological roles of these homologs are often unclear. A few studies have demonstrated the role of KaiB and KaiC homologs in stress responses in e.g. *Legionella pneumophila*^19^ and *Pseudomonas* species^20^. However, other Kai homologs are involved in the regulation of diurnal rhythms outside the cyanobacterial lineage. These include e.g. KaiB and KaiC homologs from the phototrophic bacterium *Rhodopseudomonas palustris*^21^. Recently, a KaiA-independent hourglass timer was reconstituted using *Rhodobacter sphaeroides* KaiC and KaiB homologs. *R. sphaeroides* KaiC exhibits a divergent extended C-terminus which is typically found in proteins belonging to the KaiC2 subgroup^22^. This C-terminal extension interacts with the protein, allowing for KaiA-independent phosphorylation. *R. sphaeroides* KaiB controls the phosphorylation-dephosphorylation cycle of KaiC depending on the ATP-to-ADP ratio, suggesting that metabolic changes during the day and night cycles drive this KaiBC clock^22^.

The cyanobacterium *Synechocystis* sp. PCC 6803 (hereafter *Synechocystis*) is a facultative heterotrophic cyanobacterium that, in contrast to *Synechococcus*, can utilize glucose as an energy and carbon source. *Synechocystis* encodes, in addition to the canonical *kaiAB1C1* gene cluster, two further *kaiB* homologs, named *kaiB2* and *kaiB3*, and two *kaiC* homologs, named *kaiC2*, and *kaiC3*^23^. For the *Synechocystis* KaiB3-KaiC3 timing system, Aoki and Onai suggested a function in the fine-tuning of the core oscillator KaiAB1C1 by modulating its amplitude and period^24^. This idea was supported by Wiegard *et al.,* who investigated the characteristics of the KaiC3 protein and proposed an interplay between the KaiB3-KaiC3 system and the proteins of the standard clock system^25^. Furthermore, autophosphorylation and ATPase activities of *Synechocystis* KaiC3 have been verified, suggesting that enzymatic activities might be conserved across the KaiC protein family ^25–27^. However, compared to *Synechococcus* KaiC, KaiC3 ATPase activity was reduced and lacked temperature compensation, an essential feature of true circadian oscillations^4, 25^. Recently, Zhao et al.^17^ used a luminescence gene reporter to study circadian gene expression in the *Synechocystis* wild type in comparison to mutant strains lacking each of the *kai* genes. They demonstrated that the *kaiAB1C1* and *kaiB3C3* genes are both important for circadian rhythms in *Synechocystis,* whereas *kaiC2* and *kaiB2* deletion mutants still showed rhythmic gene expression, which is in agreement with previous suggestions by Aoki and Onai^24^. Phenotypic mutant analysis by our group revealed that two systems function in the autotrophy/heterotrophy switch, especially affecting heterotrophic growth. In contrast to the study by Zhao *et al.*^17^, the deletion of *kaiC3* in the motile *Synechocystis* strain (PCC-M in^28^) used in our study had no effect on growth under light/dark cycles. However, the mutant strain displayed a growth defect under chemoheterotrophic conditions in the dark compared to the wild type^25, 29^. This impairment was less severe in comparison with the Δ*kaiAB1C1*-deficient strain, which completely lost its ability to grow in the dark. Notably, complete deletion of *kaiC2* was not possible in the wild-type strain used in our laboratory. Although Zhao et al.^17^ clearly showed that deletion of the *kaiC3* and *kaiB3* genes affects the circadian rhythm of *Synechocystis*, it remains unclear whether the KaiB3-KaiC3 system can function as an oscillator. How can such a minimal system maintain circadian rhythmicity without KaiA? *Prochlorococcus* MED4, which lacks a *kaiA* gene in the entire genome, is suggested to have no true circadian rhythmicity^30, 31^. Moreover, *Synechocystis* KaiC3 lacks the extended C-terminus, which is crucial for the oscillation of the *R. sphaeroides* KaiBC hourglass timer^22^.

In *Synechococcus*, the KaiA protein functions as a homodimer and harbors two distinct domains connected by a linker sequence^32–34^. The N-terminal domain is similar to bacterial response regulators but lacks the aspartate residue crucial for phosphorylation; hence, it is designated as a pseudoreceiver domain (PsR domain)^32^. This domain was shown to bind the oxidized form of quinones and is therefore able to directly sense the onset of darkness and forward signals to the C-terminal domain^32, 35^. The C-terminus has a four-helix bundle secondary structure and is highly conserved within Cyanobacteria. The domain harbors the KaiA dimer interface and the KaiC binding site, and is necessary to stimulate the autophosphorylation activity of KaiC^32, 34^. Mutations in *kaiA*, resulting in altered periodicity, were mapped throughout both domains, indicating their importance for rhythmicity^34, 36^.

To date, the regulatory network of the KaiB3-KaiC3 system in *Synechocystis* has remained enigmatic, as it does not interact with KaiA and does not utilize the SasA-RpaA output pathway, suggesting alternative yet unidentified components for KaiB3-KaiC3-based signal transduction^37^. In a large-scale protein-protein interaction screen, a potential interaction partner of KaiC3 was identified^38^. This protein, Sll0485, was categorized as a NarL-type response regulator and could be a potential element in the KaiB3-KaiC3 signaling pathway^39^.

In this study, we computationally characterized Sll0485 and detected strong co-occurrences of the KaiB3-KaiC3 system with Sll0485 in the genomic context of Cyanobacteria and other bacteria. Bioinformatics analysis highlighted a resemblance between the N-terminal domain of the protein and the receiver domain of NarL-type response regulators, yet the C-terminal domain shared similarities with KaiA homologs. Therefore, we investigated the effects of Sll0485 on KaiC3 phosphorylation. Sll0485 increased the phosphorylation of KaiC3 *in vitro* and *in vivo*. We observed Sll0485-dependent 24-hour oscillations of KaiC3 phosphorylation in *Synechocystis* cells grown under light/dark and continuous light conditions. Those 24h oscillations of KaiC3 phosphorylation could be reconstituted *in vitro* by incubation with Sll0485 and KaiB3. Deletion of *sll0485* led to impaired viability during mixotrophic and heterotrophic growth, in line with previous studies on the KaiB3-KaiC3 system^25^. Thus, we propose that Sll0485 is a novel KaiA-like homolog linked to the KaiB3-KaiC3 system which together with the standard KaiA1B1C1 system controls circadian rhythms and the phototrophy-to-heterotrophy switch in *Synechocystis*.

## Results

### KaiA3 is a chimeric protein harboring a NarL-type response regulator domain at the N-terminus and a conserved KaiA-like motif at the C-terminus

The canonical clock genes, *kaiABC* and *kaiA1B1C1*, form a cluster in *Synechococcus* and *Synechocystis*, respectively. In contrast, the *kaiB3* and *kaiC3* genes of *Synechocystis* are localized in different regions of the chromosome (Fig. S1A). Here, the *kaiB3* gene forms a transcriptional unit with the upstream open reading frame *sll0485.* Sll0485 has been annotated as a NarL-type response regulator^39^. Using reciprocal BLAST analyses, we detected orthologs of Sll0485 in 15 cyanobacterial species (16.5% of cyanobacterial species contained at least one KaiC homolog), mainly belonging to the order *Chroococcales*^40^ (Data S1), and in five bacterial genera outside of Cyanobacteria, namely *Roseiflexus*, *Chloroflexus*, *Chloroherpeton*, *Rhodospirillum*, and *Bradyrhizobium*.

Owing to the genetic context, we aligned the cyanobacterial Sll0485 orthologs with both, a NarL-type response regulator (Fig. S2) and cyanobacterial KaiA proteins (Fig. 1A). The canonical NarL protein consists of an N-terminal receiver domain, a linker, and a C-terminal DNA-binding domain with a helix-turn-helix motif^39, 41^. The N-terminus of the Sll0485 orthologs is conserved and indeed shows limited homology to NarL-type response regulators (Fig. S2). However, the similarities to the NarL protein decreased in the C-terminus (Fig. S2). Concurrently, conservation between Sll0845 and the KaiA protein family increased (Fig. 1A). The conserved residues in the C-terminus correspond to structurally important features of the *Synechococcus* KaiA protein, such as α-helical secondary structures, the KaiA dimer interface, or residues critical for the KaiA-KaiC interaction^32, 33^ (Fig. 1A,). Additionally, the lack of conservation in the N-terminus compared to that observed in known KaiA orthologs is consistent with the results of Dvornyk and Mei, who proposed that different N-terminal domains exist for KaiA homologs for functional diversification^42^. Because of its similarity to KaiA and synteny with the *kaiB3* gene, we named the hypothetical Sll0485 protein KaiA3. Furthermore, to facilitate the distinction of KaiA homologs, we will use the name KaiA1 for the *Synechocystis* KaiA core clock homolog Slr0756.

**Fig. 1.**
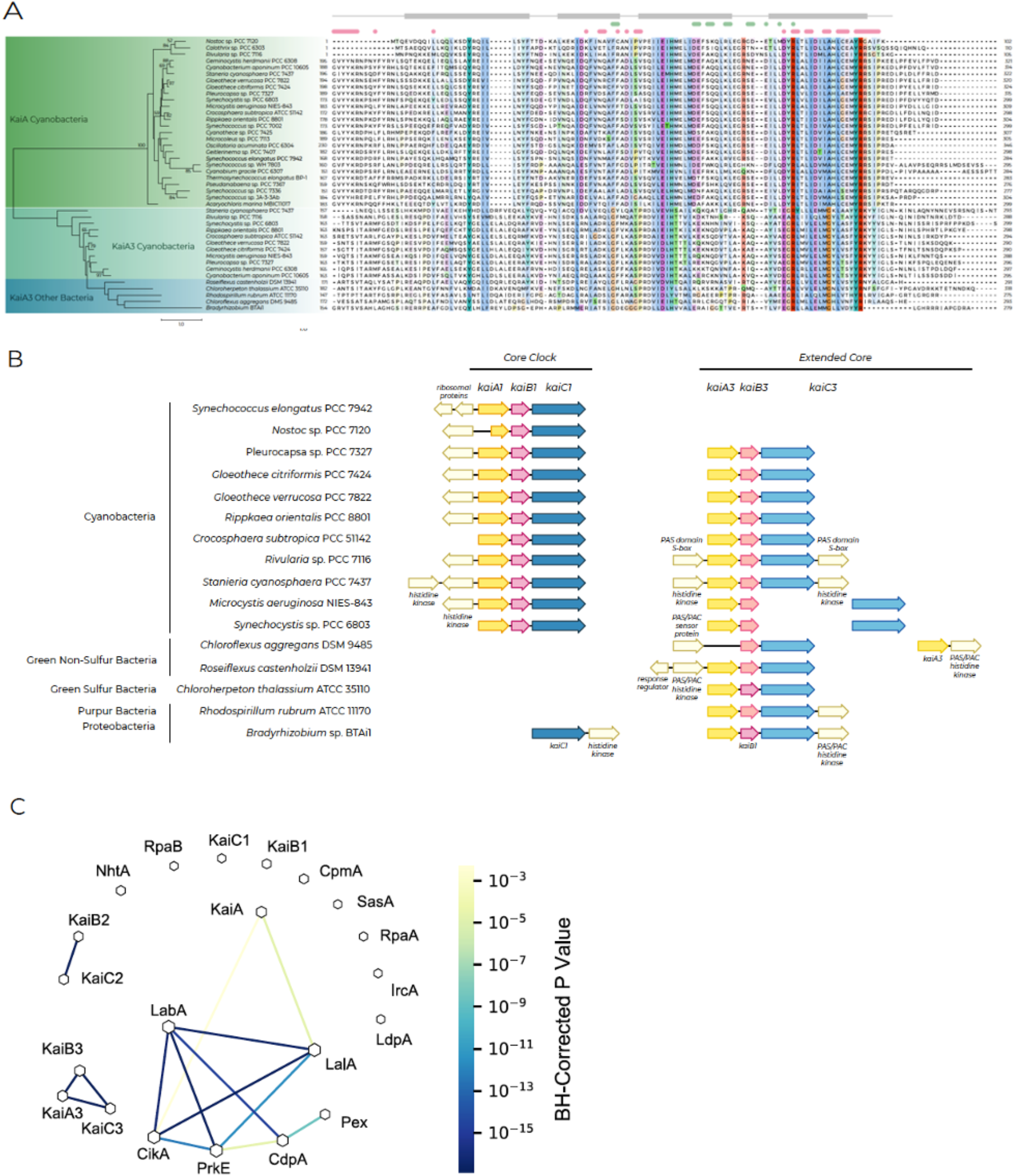
Bioinformatic analyses of Sll0485 (KaiA3). (A) Multiple sequence alignment and maximum likelihood-inferred phylogenetic reconstruction of KaiA3 and selected KaiA orthologs. The sequences were aligned with Mafft (L-INS-i default parameters, Jalview), trimmed to position 168 of the C-terminus of *Synechococcus* KaiA and are represented in the Clustalx color code with conservation visibility set to 25%. Marks above the alignment refer to *Synechococcus* KaiA as a reference. Light green bars and dots indicate residues critical for KaiC interaction, light pink bars and dots represent residues important for dimerization, and light gray blocks outline residues forming α-helices as secondary structures. Aligned sequences were used to infer a maximum likelihood protein tree. The scale bar indicates one substitution per position. Bootstrap values (n=1000) are displayed on the branches. Bootstrap values less than 50 are not shown. (B) Synteny analysis of *kaiA1B1C1* compared to *kaiA3*, *kaiB3*, and *kaiC3* genes for selected bacterial species. Analysis was performed with the online tool SyntTax, a prokaryotic synteny and taxonomy explorer (https://archaea.i2bc.paris-saclay.fr/synttax/; 2020-06-08). Default settings were used for analysis (best match, 10% norm. Blast). (C) Co-occurrence of KaiA3 using pairwise Fisher’s exact test with circadian clock proteins in Cyanobacteria. Network of significant co-occurring circadian clock factors from Schmelling *et al.*^26^, including KaiA3 in Cyanobacteria. The line color corresponds to the level of significance resulting from pairwise Fisher’s exact test. Missing links were those with a p-value higher than 0.01. The node size is proportional to the degree of that node.

The gene tree resulting from the multiple sequence alignment (Fig. 1A) distinctly separated KaiA3 from canonical KaiA orthologs. To further investigate the evolutionary relationship of KaiA3, multiple sequence alignments of the C-termini of orthologs of KaiA3, KaiA, and Slr1783 (Rre1) as a reference for NarL orthologs in Cyanobacteria^43^ were used to construct a phylogenetic tree (Fig. S3). Here, KaiA3 orthologs form a distinct clade at the basis of the KaiA orthologs when compared to both orthologous groups of Slr1783 (Rre1)/NarL (*E. coli*, UniProtKB - P0AF28) and KaiA simultaneously (Fig. S3). In summary, these findings strengthen the idea that the C-terminus of KaiA3 functions similarly to that of KaiA.

We further constructed three-dimensional models of KaiA3 to gain a better understanding of its potential functions. To date, no structure is available for KaiA3, and it is impossible to generate a reliable three-dimensional model covering the full-length KaiA3 sequence because of the enigmatic structure of the linker region, for which no significant similarities could be detected. However, secondary structure prediction suggested that the N-terminus structurally aligns with NarL (Fig. S4A). Therefore, we modeled the N-terminus (residues 1-140) and the remaining part of the sequence separately (residues 141-299). For the N-terminus, numerous hits for response regulator domains were obtained, with *E. coli* NarL (PDB 1A04) showing the highest degree of sequence similarity. The 3D-model structures of KaiA3 are highly similar and display the canonical fold of response regulator domains: a central five-stranded parallel β-sheet flanked on both faces by five amphipathic α-helices and a phosphorylatable aspartate residue in the β3-strand (Fig. S4B). This aspartate residue (D65) plays a role in response regulator phosphorylation (Fig. S2, blue stars) and is conserved in all species, except *Pleurocapsa* and *Microcystis*. Thus, most KaiA3 homologs, including the *Synechocystis* protein, harbor a potential phosphorylation site. Furthermore, the structure superimposes well on the PsR domain of KaiA, even though the PsR domain lacks the phosphate-accepting aspartate residue and the α4-helix between the β4- and β5-strands (Fig. S4B). The amino acid sequence between the β4- and β5-strands shows the least conservation between KaiA and KaiA3, yet the level of sequence conservation in this region is generally low for KaiA and its homologs^34^. In contrast to the N-terminal response regulator domain, the C-terminal domain of KaiA3 revealed a unique fold, which has only been detected in KaiA thus far^44^, and the N-terminal domain of the phosphoserine phosphatase RsbU from *Bacillus subtilis*^45^, namely, a unique four α-helix bundle constituting the KaiA-like motif (Fig. S4C). In conclusion, we propose that KaiA3 consists of two protein modules: i) the N-terminal domain, resembling a NarL-type response regulator receiver domain, including its phosphorylation site, and ii) the C-terminal domain displaying features of a KaiA-like motif. This is particularly intriguing because putative *kaiA* orthologs outside Cyanobacteria have not been identified until recently^42^.

### Conserved synteny and co-occurrence of KaiA3 and the KaiB3-KaiC3 system among prokaryotes

As in *Synechocystis*, we found the *kaiA3* gene upstream of *kaiB3* in all the analyzed cyanobacterial genomes. Furthermore, the *kaiA3B3* cluster is usually extended by *kaiC3*, with only two exceptions (*Synechocystis* and *Microcystis aeruginosa* NIES-843), which resemble the structure of the canonical *kaiABC* gene cluster (Fig. 1B). Interestingly, *kaiA3B3C3* synteny was also found in other prokaryotic genomes that harbor orthologs of *kaiA3*, except for *Chloroflexus aggregans* DMS 9485 (Fig. 1B). Furthermore, we detected strong significant co-occurrences between KaiA3 and KaiB3 (p<0.0001) as well as between KaiA3 and KaiC3 (p<0.0001; Fig. 1C) in organisms encoding KaiC1. The co-occurrence of KaiB3 and KaiC3 has been previously shown^26^. Thus, KaiA3 forms a distinct set of proteins with KaiB3 and KaiC3, which show no further significant co-occurrence with other clock components (Fig. 1C,^26^). Altogether, both datasets suggest a functional relationship between KaiA3 and the KaiB3-KaiC3 system.

### KaiA3 interacts with and promotes autokinase activity of KaiC3

Using yeast two-hybrid (YTH) experiments, we verified the interaction between the clock proteins KaiC3 and KaiA3 (Fig. 2A, Fig. S5), consistent with a previous large-scale protein-protein interaction analysis by Sato *et al*.^38^. Although KaiA3 clearly interacted with KaiC3, an interaction with KaiB3, the second element of the KaiB3-KaiC3 clock system, was not detected (Fig. S5B). This is not surprising, as it has been demonstrated that the interaction of the *Synechococcus* proteins KaiA and KaiB requires the presence of KaiC^46^. To further characterize the interaction of the proteins *in vitro*, we heterologously expressed different Kai proteins in *E. coli* and analyzed complex formation using clear-native PAGE (Fig. 2B and Fig. S6). The His-tagged KaiA3 protein (monomer: 35 kDa) migrated as a single band approximately 100 kDa in size, indicating the formation of KaiA3 homo-oligomers, at least dimers. *Synechococcus* KaiA migrated at ∼60 kDa, in line with previous results^47^, confirming the formation of KaiA dimers. The discrepancy in the migration pattern between KaiA3 (His-tagged) and KaiA (GST-tag removed) might be due to differences in their predicted charge (-19.17 for KaiA and -7.94 for KaiA3, respectively, at pH 7.0). Recombinant KaiB3 (monomer: 12 kDa) was shown to form monomers and tetramers after size exclusion chromatography^25^. KaiB3 displayed three distinct bands in the native gels (Fig. 2B). The two lower bands most likely represent the monomeric and tetrameric forms, whereas the uppermost band (∼67 kDa) could be an impurity in the protein preparation. Recombinant KaiC3 was produced with an N-terminal Strep-tag^25^. Strep-tagged KaiC3 (monomer: 58 kDa) migrated as one band between 272 and 450 kDa and could represent a hexameric complex (348 kDa). Incubation of KaiC3 with KaiA3 alone led to protein accumulation in the wells in native PAGE, indicating precipitation of the KaiA3/KaiC3 complex in the absence of KaiB3 (Fig. 2B). However, the interaction between KaiA3 and KaiC3 was validated by immunoprecipitation-coupled liquid chromatography-mass spectrometry (LC-MS) analysis of FLAG-tagged KaiC3 (Fig. S7). Furthermore, the experiments did not reveal any interactions between KaiA3 and either KaiC1 or KaiC2 (Fig. S5, Fig. S7), indicating specificity of the KaiA3-KaiC3 interaction. No complex formation was detected between KaiA3 and KaiB3 (Fig. 2B, Fig. S5 and Fig. S6). In contrast, the formation of a large protein complex was observed when all three clock components, KaiA3, KaiB3, and KaiC3, were incubated together for 16 h at 30°C (Fig. 2B; Fig. S6). The size matches that of a complex consisting of one KaiC3 hexamer, six KaiA3 dimers, and six KaiB3 monomers (840 kDa). The presence of KaiA3 in the complex was validated by western blot analysis using an anti-His antibody (Fig. 2B, Fig. S6). As expected, no such complex was formed when KaiA3 was replaced with *Synechococcus* KaiA (Fig. 2B). Moreover, no such complex was formed when KaiB3 was replaced by its isoform KaiB1, suggesting that KaiB3 is specific for KaiA3 as well and that KaiB3 might recruit KaiA3 to the KaiC3/KaiB3 complex (Fig. S6).

**Fig. 2.**
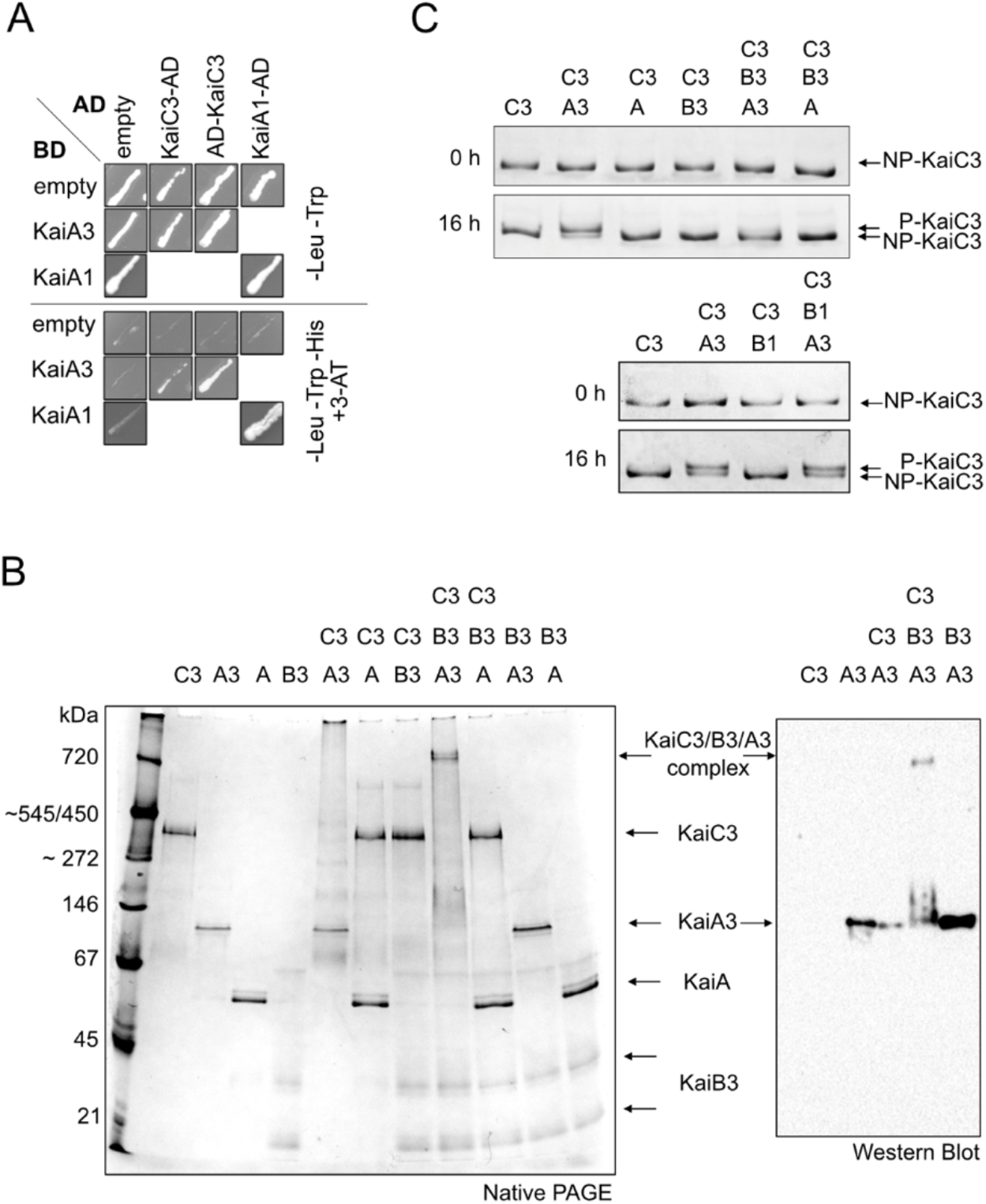
Analysis of KaiA3 protein interactions and KaiC3 phosphorylation. (A) YTH interaction analysis of KaiA3 with KaiC3. The KaiA1 dimer interaction was used as a positive control. YTH reporter strains carrying the respective bait and prey plasmids were selected by plating on complete supplement medium (CSM) lacking leucine and tryptophan (-Leu -Trp). AD, GAL4 activation domain; BD, GAL4 DNA-binding domain; empty, bait, and prey plasmids without protein sequence (only AD/BD domain). The physical interaction between bait and prey fusion proteins was determined by growth on complete medium lacking leucine, tryptophan, and histidine (-Leu -Trp -His) and the addition of 12.5 mM 3-amino-1,2,4-triazole (3-AT). The BD was fused to the N-terminus of KaiA3. For a clear presentation, spots were assembled from several replicate assays (original scans are shown in Fig. S5). (B) Interaction analysis of the recombinant Kai proteins on native polyacrylamide gels. Proteins were incubated for 16 h at 30°C and subsequently subjected to 4-16% clear native PAGE. Gels were either stained with Coomassie Blue (left side) or blotted and immunodecorated with a monoclonal anti-His antibody to detect recombinant KaiA3-His6 (right side). Recombinant *Synechococcus* KaiA was used for comparison. (C) KaiC3 phosphorylation depends on the presence of KaiA3 and KaiB3. KaiC3 was dephosphorylated by incubating for 18 h at 30°C prior to the start of the assay (NP-KaiC3). 0.2 µg/µl NP-KaiC3 was incubated at 30°C in the presence or absence of 0.1 µg/µl *Synechocystis* KaiA3, KaiB3 and KaiB1 and *Synechococcus* KaiA, respectively. Aliquots were taken at 0 h and 16 h, followed by separation on a high-resolution LowC SDS-PAGE gel in Tris-Tricine buffer and staining with Coomassie blue. A slow-migrating band representing the phosphorylated form of KaiC3 (P-KaiC3) was observed only in the presence of KaiA3.

Previous studies have shown that KaiC3 has autokinase activity, which is independent of KaiA1^25, 27^. Since our studies revealed an interaction between KaiC3 and KaiA3, we were interested in probing the influence of KaiA3 on the phosphorylation of KaiC3. The recombinant Kai proteins described above were used for this purpose. KaiC3 was incubated for 16 h at 30°C in the presence or absence of other Kai proteins, and its phosphorylation state was analyzed by SDS-PAGE (Fig. 2C), and LC-MS/MS (Fig. S8). Since KaiC3 was partially phosphorylated after purification from *E. coli,* the protein preparation was incubated for 18 h at 30°C prior to the start of the assays. During this incubation period, KaiC3 autodephosphorylated, as is typical for KaiC proteins (Fig. 2C, NP-KaiC3)^44^. Addition of KaiA3 led to phosphorylation of KaiC3, while the presence of *Synechococcus* KaiA had no influence on the phosphorylation state of KaiC3. In contrast, KaiC3 dephosphorylation was enhanced by KaiB3 (Fig. 2C, upper panel). Replacing KaiB3 with its isoform, KaiB1, in samples containing KaiA3, maintained KaiC3 in the phosphorylated state (Fig. 2C, lower panel). Analysis of KaiC3 phosphorylation by LC-MS/MS-identified the neighboring residues Ser423 and Thr424 as phosphorylation sites, which are conserved across KaiC homologs (Fig. S8). Based on these analyses, we conclude that KaiA3 likely has a KaiA-like function in promoting the phosphorylation of KaiC3. Neither *Synechococcus* KaiA nor *Synechocystis* KaiB1 could substitute for KaiA3 or KaiB3, respectively, demonstrating that the *Synechocystis* KaiA3/KaiB3/KaiC3 proteins represent a separate functional complex. Only KaiA3 stimulated the autokinase activity of KaiC3, which in turn promoted its interaction with KaiB3. Interaction with KaiB3, but not KaiB1, enhances the dephosphorylation of KaiC3.

### KaiC3 phosphorylation oscillates in vitro and in Synechocystis cells

The opposing effects of KaiA3 and KaiB3 on KaiC3 phosphorylation imply that these three *Synechocystis* proteins may form a functional *in vitro* oscillator. We monitored the phosphorylation of KaiC3 in concert with KaiB3 and various concentrations of KaiA3 over a period of 48 h (Fig. 3A, B; Fig. S9). In the presence of 1.4 µM and 2.8 µM KaiA3 (corresponding to a ∼1:1.2 and 1:2.4 stoichiometry of KaiA3:KaiC3), we could reconstitute ∼24h oscillations in KaiC3 phosphorylation (Fig. 3A, B; Fig. S9). Compared to the *Synechococcus* KaiABC oscillator^48, 49^, lower KaiA3 concentrations failed to generate oscillations and the protein was mainly dephosphorylated. The stimulating effect of KaiA3 on KaiC3 phosphorylation was saturated at a KaiA3 concentration of 4.2 µM, which corresponds to a KaiA3:KaiC3 stoichiometry of 1:0.8. Hence, the KaiC3 oscillations were clearly dependent on the KaiA3 concentration.

**Fig. 3.**
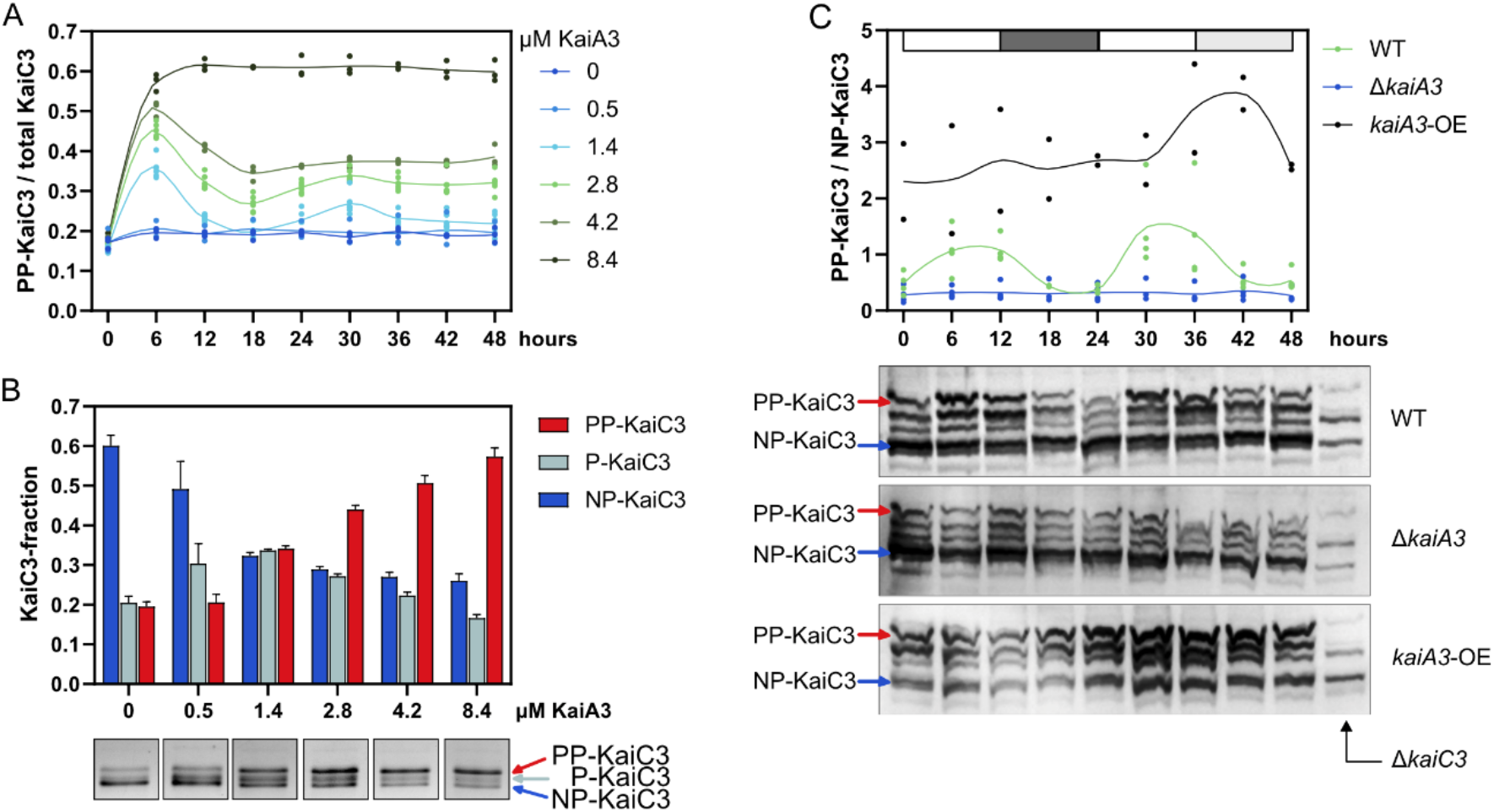
Analysis of KaiA3-dependent KaiC3 phosphorylation. (A) KaiC3 (3.4 µM) was incubated with KaiB3 (7.4 µM) and various concentrations of KaiA3 at 30°C. Aliquots incubated for the indicated time periods were applied to a high-resolution LowC SDS-PAGE gel, proteins were separated in Tris-glycine buffer, and the relative band densities of the different KaiC3 phosphorylation states: unphosphorylated (NP), single-phosphorylated (P), and double-phosphorylated (PP) were estimated densitometrically. (A) *In vitro* ratio of fully phosphorylated KaiC3 (PP-KaiC3) to total KaiC3 at various concentrations of KaiA3. Dots display replicates (n=3); the line represents an akima spline curve. Assays with 1.4 µM KaiA3 and 2.8 µM KaiA3 were each analyzed twice on the gel, resulting in 6 replicates in total. Representative gels from each assay are shown in Fig. S9A. (B) Detailed analysis of KaiC3 phosphorylation after 6h of incubation with different KaiA3 concentrations. The fractions of double (PP), single (P), and unphosphorylated KaiC (NP-KaiC3) are plotted as average +SD from the three assays, also shown in A. Below the graph, representative band patterns are shown (assembled from Fig. S9A). (C) The ratio of fully phosphorylated KaiC3 (PP-KaiC3) to non-phosphorylated KaiC3 (NP-KaiC3) in *Synechocystis* wild-type, *kaiA3* mutant (τι*kaiA3*), and *kaiA3* overexpression (*kaiA3-OE*) strains (Fig. S1). Whole cell extracts were separated using Phos-tag SDS-PAGE and immunodecorated with a KaiC3-specific antiserum. Samples were collected every 6 h from cells grown in a 12-h light/dark cycle, followed by constant light. The white and dark gray boxes represent light and dark periods, respectively, and the light gray box represents the subjective night. Representative blots are shown. Whole cell extracts from the *Synechocystis ΔkaiC3* mutant (12 h time point) were used as a control. Dots in the graph display the replicates (n=2-3); the line represents an akima spline curve. Plots were generated using GraphPad Prism, version 9.5.1.

To evaluate whether the self-sustained KaiC3 phosphorylation rhythms detected above are also present in *Synechocystis* cells and are diurnal or circadian in nature, we grew cells in a light/dark cycle, followed by constant illumination. We separated whole-cell extracts on a Phostag gel and identified KaiC3 by Western blot analysis (Fig. 3C) using a KaiC3-specific antibody^27^. We detected 4-5 bands which partially overlapped or were slightly shifted in comparison to the bands detected in the Δ*kaiC3* strain. It seems that there is some cross-reaction with KaiC1, KaiC2 or another protein The two prominent bands indicated in Fig. 3C and which are absent in the Δ*kaiC3* strain most probably reflect two different phosphorylation states of KaiC3. Based on the in vitro data with the isolated Kai proteins and their similar migration patterns in Phos-tag SDS-PAGE analysis compared to the whole cell extract (Fig. S9C), we suppose that the very upper (red arrow) and one of the lower bands (blue arrow) in Fig. 3C represent the fully phosphorylated and non-phosphorylated forms of KaiC3, respectively. In the Δ*kaiA3* mutant, the lowest band was mainly present, indicating that KaiC3 was mostly dephosphorylated in this strain. Incubation of KaiC3 with Lambda phosphatase resulted in comparable accumulation of the lower band (Fig. S9D). In contrast, in the KaiA3 overexpression strain, KaiC3 was highly phosphorylated in comparison to the wild type (Fig. 3C). In addition, two or more bands were detected in the in vitro assays, as well as in the cell extracts (Fig. 3B, C; Fig. S9C) which partly overlapped with an unspecific band detected in the Δ*kaiC3* strain in Phos-tag SDS-PAGE analysis. These bands might reflect single phosphorylated states of KaiC3. In summary, our in vitro and in vivo data demonstrate that KaiC3 phosphorylation strongly depended on KaiA3. Furthermore, KaiC3 phosphorylation showed sustained oscillations with a 24 hours rhythm in *Synechocystis*, hence displaying a characteristic feature of a circadian oscillator.

### Deletion of kaiA3 impacts growth and viability during mixotrophic and chemoheterotrophic growth

What is the function of this additional Kai protein oscillator in *Synechocystis*? In our laboratory, deletion of *kaiC3* led to growth impairment in complete darkness on glucose, but not in light/dark cycles^25^; thus, the *kaiA3* knockout mutant (Δ*kaiA3*) was analyzed under various growth conditions. The cells were grown in liquid culture under constant light, plated on agar at different dilutions, and grown photoautotrophically (Fig. 4A) and photomixotrophically (Fig. 4B) under continuous light and 12-h light/12-h dark cycles or chemoheterotrophically (Fig. 4C). Because the strains grew very slowly under chemoheterotrophic conditions, the cells were spotted at higher concentrations under these conditions. There were no differences in the viability of the mutant strains in comparison to that of the wild type under photoautotrophic conditions in continuous light and light/dark cycles. Under photomixotrophic conditions, the Δ*kaiA3* strain showed less viability, which was partly restored by re-insertion of *kaiA3*. It appears that the amount of KaiA3 is critical for the function of the system, which is consistent with our data on KaiA3-dependent KaiC3 phosphorylation (Fig. 3). Surprisingly, the mutant strain lacking all three alternative *kai* genes (Δ*kaiA3B3C3)* exhibited a different growth phenotype under photomixotrophic conditions. In light/dark cycles, this strain grew well and seemed to have some advantages in comparison to the wild type (Fig. 4B). Spot assays under chemoheterotrophic conditions provided a clearer picture; the mutant strain lacking *kaiA3* and the triple knockout showed a similar phenotype. They were unable to grow in complete darkness, and this ability was fully restored in the complementation strain (Fig. 4C). These results coincide with previously detected impairments displayed by the Δ*kaiC3* strain during chemoheterotrophic growth^25^, strengthening the idea that the non-standard KaiA3-KaiB3-KaiC3 system is a regulatory complex with the same function.

**Fig. 4.**
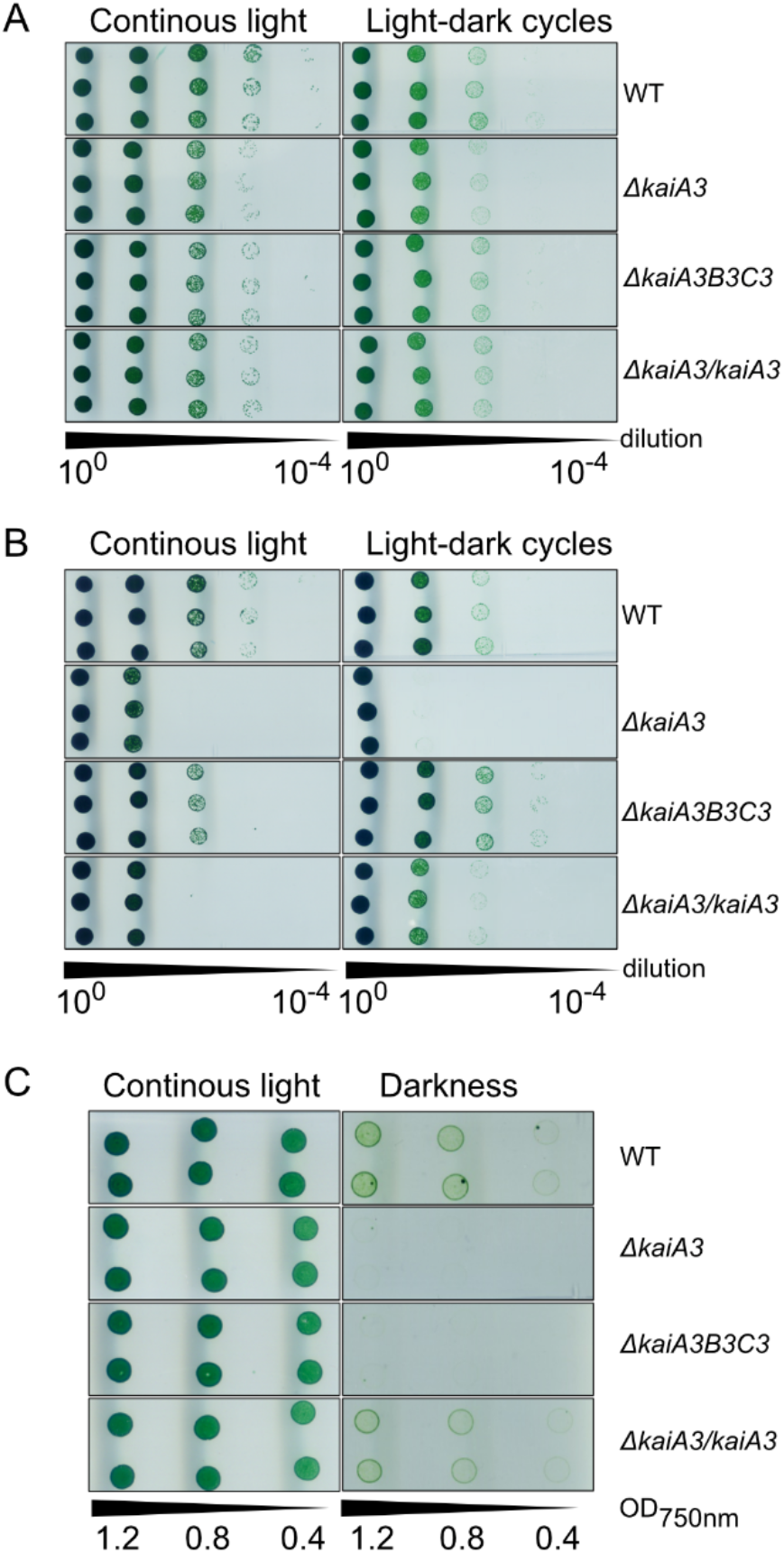
Deletion of *kaiA3* results in growth defects during mixotrophic and chemoheterotrophic growth. Proliferation of the wild type (WT), the Δ*kaiA3* and Δ*kaiA3B3C3* deletion mutants, and the Δ*kaiA3*/*kaiA3* complementation strain was tested under (A) phototrophic (continuous light, - glucose), (B) photomixotrophic (continuous light, + glucose), and (C) heterotrophic (darkness, + glucose) conditions. Strains were grown in liquid culture under constant light, and different dilutions were spotted on agar plates and incubated under the indicated light conditions with a light phase corresponding to 75 µmol photons m^-2^ s^-1^ white light. Representative result from three independent experiments are shown. (A) Cultures were diluted to an OD_750nm_ value of 0.4, and tenfold dilution series were spotted on agar plates. Plates were analyzed after 6 or 8 days of continuous light and 12h/12h light/dark cycles, respectively. (B) Same as (A), but the cells were spotted on agar plates containing 0.2 % glucose. (C) Cultures were diluted to OD_750nm_ values of 1.2, 0.8, and 0.4, and spotted on agar plates supplemented with 0.2% glucose. The plates were analyzed after 3 and 26 d of continuous light and darkness, respectively.

## Discussion

Our knowledge of the function, composition, and network of clock systems in prokaryotes, including cyanobacteria, is increasing steadily. Even though multiple copies of the core clock proteins KaiB and KaiC are encoded in bacterial genomes, the canonical KaiA was found only as a single copy in Cyanobacteria yet^26, 27, 42, 50^. By identifying a chimeric KaiA3 and verifying its interaction with the KaiB3-KaiC3 complex, we added another component to the diversity of bacterial clock systems.

### KaiA-like proteins outside of Cyanobacteria and primordial clocks

In addition to KaiA3, new putative KaiA orthologs have been bioinformatically identified in prokaryotes outside Cyanobacteria^42^. Therefore, we suggest that such proteins may play a previously overlooked role in KaiB-KaiC-based systems. Exploring this possibility could provide valuable insights into unanswered research questions, such as the mechanism responsible for the rhythmic processes observed in *Rhodospirillum rubrum.* Indeed, this purple bacterium lacks KaiB1 and KaiC1 orthologs but possesses KaiA3, KaiB3, and KaiC3 (53) (Figure 1). Notably, the recently described oscillator from *Rhodobacter sphaeroides* (*Rhodobacter*), which consists of homologs of KaiC2 and KaiB2, can form an hourglass timer. This primordal *Rhodobacter* clock can function without KaiA. However, the *Rhodobacter* KaiB2-KaiC2 system requires an environmental signal to reset the clock^22^. A similar primordial clock has been suggested to be present in *Rhodopseudomonas palustris*^21^ and the cyanobacterium *Prochlorococcus* MED4^30, 31^. However, other bacterial KaiB and KaiC homologs, including the KaiC2-KaiB2 system from *Synechocystis*, are believed to have clock-independent functions^17, 51, 52^.

It has been proposed that *kaiC* is the oldest evolutionary member of circadian clock genes^50^. KaiC homologs can be found even in Archaea where it was found to control e. g. motility of *Sulfolobus acidocaldarius* by protein interaction^53^. The later addition of KaiB was enough to form a primordial timekeeper which needs a signal for daily resetting of the clock^21, 22, 30, 31^. In *Rhodobacter* KaiC2, dephosphorylation is regulated by the stability of coiled-coil interactions between two connected hexamers as well as by KaiB^22^. However, whether autophosphorylation or dephosphorylation dominates depends primarily on the ATP/ADP ratio. Hence, the KaiC2-KaiB2 timer cannot oscillate autonomously but responds to changing ATP/ADP levels. Therefore, it was suggested that the *Rhodobacter* clock represents an ancient timer that depends on changes in photosynthetic activity during the day-night switch^22^.

With the evolution of KaiA, a self-sustained oscillator was developed that allowed for true circadian oscillations in gene expression, which can be observed in cyanobacteria. Why does KaiC require KaiA to drive persistent oscillations? By default, the A-loops of *Synechococcus* KaiC hexamers adopt a buried conformation, which inhibits autophosphorylation. Only the binding of KaiA favors phosphorylation by stabilizing A-loop exposure^5^. In contrast, *Rhodobacter* KaiC2 constantly exposes its A-loops, sterically allowing high intrinsic phosphorylation^22^.

The interacting residues between KaiA and KaiC are less conserved in both *Synechocystis* KaiA3 and KaiC3^27, 54^ (Fig. 1). Since we demonstrated an interaction between KaiC3 and KaiA3, it is likely that co-evolution of the two proteins occurred. Another remarkable feature of *Rhodobacter* KaiC2 is that the latter displays an extended C-terminus that connects two hexamers via coiled-coil interactions to adopt a homododecamer instead of a typical hexamer^22, 55^. KaiC3 does not have such an extended C-terminus^27^, and we only observed the formation of hexamers or smaller oligomers^25^ (Fig. 2).

### The two-domain architecture of KaiA3 and complex formation

KaiA3 formed a distinct clade at the basis of the KaiA clade. Apart from its presence in the N-terminal domain of phosphatase RsbU of *Bacillus subtilis*, a distinctive structure of the KaiA C-terminus has rarely been observed^45^. RsbU acts as a positive regulator of the alternative sigma factor B, which is involved in the general stress response^56^. The N-terminal domain of RsbU forms dimers similar to KaiA, and the proposed binding site for its corresponding activator, RsbT, is in an equivalent location to the KaiC-binding site on KaiA^45^. These findings may reflect how protein domains change during evolution, while their original functions are conserved. However, a link between RsbU and the recently proposed circadian clock in *Bacillus subtilis* has not yet been identified^57^. Moreover, circadian rhythms have been observed in several prokaryotes that do not encode Kai orthologs, suggesting the convergent evolution of circadian rhythms in prokaryotes^57, 58^. Further in-depth analyses are needed to elucidate whether KaiA3, together with KaiB3 and KaiC3, or the well-studied *Synechococcus* circadian clock present a more ancestral system, because analysis of a larger dataset recently suggested that the canonical *kaiA* gene evolved at the same time as cyanobacteria^42^.

Taken together, these data are consistent with a model in which KaiA3 can fulfill the functions of a true KaiA homolog, such as dimerization, binding to KaiC3, and enhancing KaiC3 autophosphorylation. Other mechanistic processes, such as sequestration to the CI ring by binding to KaiB3, remain to be investigated but are clearly possible. By mixing KaiA3, KaiB3, and KaiC3, we reconstituted a *bona fide in vitro* oscillator (Fig 3A), suggesting that the observed in vivo oscillation of KaiC3 phosphorylation can run independently of the KaiA1B1C1 clock, and that the amount of KaiA3 is critical for the phosphorylation rhythm.

The *Rhodobacter* hourglass-like timer requires environmental cues for daily resetting. However, entrainment by metabolites has also been described for more elaborate, true circadian oscillators. In addition to entrainment by the input kinase CikA^59^, the *Synechococcus* clock can be entrained directly by the ATP/ADP ratio and oxidized quinones^35, 60^. Moreover, CikA does not sense light directly but perceives the redox state of the plastoquinone pool^61, 62^. In addition, glucose feeding can entrain *Synechococcus* when engineered to take up glucose^63^. In plants, it has been demonstrated that both exogenous sugars and internal sugar rhythms resulting from cyclic photosynthetic activity entrain the clock^64^. *Synechocystis* can naturally utilize glucose, which may make it even more susceptible to metabolic entrainment by sugars. Notably, the *Synechocystis* wild-type strain used in this study was able to grow in complete darkness when supplemented with glucose. This is different from an earlier study that showed that *Synechocystis* needs a 5 min blue-light pulse at least once a day to grow heterotrophically in the dark^65^. The authors described this behavior as light-activated heterotrophic growth. There are no studies that explain why cells require this short light pulse, but it is also clear that our laboratory strain grows fully chemoheterotrophically^29^.

In contrast to *Synechococcus*, CikA from *Synechocystis* is a true photoreceptor that binds a chromophore^66^. Thus, it remains unclear whether CikA has a similar function in both cyanobacteria, and whether it interacts with both circadian clock systems in *Synechocystis*. The high structural similarity of the N-terminal domain of KaiA3 to response regulator domains from other organisms indicates that the core structure and activity are maintained, while adaptivity and variation provide specificity for acting in distinct pathways^24^. Within KaiA3, the aspartate residue crucial for phosphorylation is conserved. Theoretically, the protein could receive an input signal from a cognate histidine kinase, which has not yet been identified. Thus, there are potentially important differences related to input and output factors, and possibly entrainment of different cyanobacterial circadian clock systems.

### The function of KaiA3 in Synechocystis

The physiological function of the KaiA3-KaiB3-KaiC3 clock system seems to be related to the different metabolic modes of *Synechocystis.* Mutants deficient in *kaiA3* lose the ability to grow chemoheterotrophically on glucose, which is an aggravated effect compared to *kaiC3*-deficient mutants, which merely show reduced growth rates during heterotrophy^25^. Similarly, in *Synechococcus,* the disruption of *kaiA* led to one of the most severe effects on activity loss and was traced back to the unbalanced output signaling of the circadian clock^67^. The overaccumulation of KaiA3 also appeared to disturb the system (Fig. S10). Such an effect was also shown for the *Synechococcus* clock system, in which increased KaiA levels promote the hyperphosphorylation of KaiC^6, 68^, thereby deactivating rhythmic gene expression^69^. Surprisingly, inactivation of the complete KaiA3-KaiB3-KaiC3 system resulted in a different phenotype. While growth in darkness on glucose was strongly affected, similar to the single mutants, photomixotrophic growth was even slightly better in the *kaiA3B3C3* strain compared to the wild type. It is possible that in the absence of KaiA3, the altered interaction of KaiC3 with KaiC1 leads to aggravated growth defects in the Δ*kaiA3* mutant. However, when a complete oscillator is missing, the KaiA1B1C1 oscillator can compensate for this under certain growth conditions.

In *Synechocystis*, Δ*kaiA3*-like phenotypes, such as impaired viability during light/dark cycles or complete loss of chemoheterotrophic growth on glucose, were also observed for Δ*kaiA1B1C1*, Δ*sasA*, and Δ*rpaA* mutants^29, 70^. For Δ*sasA,* it was shown that the mutant strain was able to accumulate glycogen but was unable to utilize the storage compound to grow heterotrophically, probably because of its inability to catabolize glucose^70^. A recent metabolomics study suggested that the growth inhibition of Δ*kaiA1B1C1* and Δ*rpaA* mutants in a light/dark cycle might be at least partly related to a defect in the inhibition of the RuBisCo enzyme in the dark and increased photorespiration, leading to the accumulation of the potentially toxic product 2-phosphoglycolate^71^. This previous study also revealed an enhanced growth defect in Δ*kaiA1B1C1* and Δ*rpaA* mutants under photomixotrophic conditions in light/dark cycles, similar to the Δ*kaiA3* strain in the current study. This further supports the idea that one of the functions of the KaiA3-B3-C3 system is to fine-tune the core clock system, KaiA1B1C1. Clearly, there is a difference in the phenotypes between our study and the results demonstrated by Zhao et al.^17^, who analyzed single and double *kaiB3* and *kaiC3* knockout strains. In light/dark cycles, the *kaiB3C3* knockout strain showed a reduced growth rate compared to the wild-type control under photoautotrophic conditions. However, under constant light, this mutant showed a reduced growth rate and was outcompeted by the wild-type cells in mixed cultures. Photoheterotrophic and heterotrophic conditions were not tested in this study. *Synechocystis* strains used in different laboratories can vary in their genome and phenotypic characteristics, including glucose sensitivity (see for example^28, 72^). As the input and output pathways of the new oscillator are unknown, it is possible that mutations in different wild-type variants lead to variations in the expression of phenotypic effects in the clock mutants.

Here, we demonstrate that KaiA3 is a novel KaiA homolog and element of the KaiC3-based signaling pathway and has canonical KaiA functions. The N-terminal half of KaiA3 may still have a response regulatory function; however, the exact mechanism remains unclear. Among other actions, KaiA3 must be placed within the regulatory and metabolic networks of *Synechocystis*. Finally, our findings in the cyanobacterium *Synechocystis* demonstrated the parallel presence of two circadian protein oscillators within a single cell.

## Materials and Methods

### Reciprocal BLAST of Sll0485 (KaiA3) and Slr1783 (Rre1)

Reciprocal BLAST was performed as described by Schmelling *et al*.^26^. The 2017 database was used for comparison with existing data on other circadian clock proteins. The protein sequences of Sll0485 (KaiA3) and Slr1783 (Rre1), as a reference for NarL response regulators^43^ from *Synechocystis,* were used as query sequences for this reciprocal BLAST.

### Co-occurrence analysis

The co-occurrence of KaiA3 with other circadian clock proteins in Cyanobacteria containing KaiC1 was examined according to Schmelling *et al.*^26^. A right-sided Fisher’s exact test was used^73^. P-values were corrected for multiple testing after Benjamini-Hochberg^74^, with an excepted false discovery rate of 10^−2^. All proteins were clustered according to their corrected p-values.

### Synteny analyses using SyntTax

The conservation of gene order was analyzed using the web tool ‘SyntTax’^75^; https://pubmed.ncbi.nlm.nih.gov/23323735/. If not mentioned otherwise, default settings (Best match, 10 % norm. BLAST) were applied. Chromosomes were selected manually according to the results of Schmelling *et al*.^26^.

### Multiple sequence alignments with Mafft and Jalview

Sequence alignments, visualization, and analysis were performed with ‘Jalview’^76^. The sequences were aligned with Mafft, and if not mentioned otherwise, default settings (L-INS-i, pairwise alignment computation method - localpair using Smith-Waterman algorithm, gap opening penalty: 1.53, gap opening penalty at local pairwise alignment: - 2.00, group-to-group gap extension penalty: 0.123, matrix: BLOSUM62) were applied^77^. For analyses of the C-terminus, the alignments were trimmed to position 168 in the KaiA reference sequence of *Synechococcus*. After trimming, the alignment was recalculated with Mafft using the aforementioned default parameters.

### 2D and 3D structure predictions

The alignments generated in Jalview were then used with ‘Ali2D’ for secondary structure prediction^78^ https://toolkit.tuebingen.mpg.de). The identity cut-off to invoke a new PSIPRED run was set to 30%. Three-dimensional protein structures were modeled using either Phyre2 or SWISS-MODEL^79, 80^

(http://www.sbg.bio.ic.ac.uk/phyre2/html/page.cgi?id=index; https://swissmodel.expasy.org/). The resulting structures were analyzed and illustrated using UCSF Chimera^81^ (https://www.cgl.ucsf.edu/chimera/).

### Phylogenetic reconstruction of protein trees

Phylogenetic reconstruction of the protein trees of Sll0485 (KaiA3), Slr1783 (Rre1)/NarL (*E. coli*, UniProtKB - P0AF28), and KaiA was achieved with MEGA X^82, 83^ using the above constructed alignments. For all alignments, a neighbor-joining tree and maximum likelihood tree were constructed and compared. To construct neighbor-joining trees, 1000 bootstrap iterations with a p-distance substitution model and a gamma distribution with three gamma parameters were used. To construct maximum likelihood trees, an initial tree was constructed using the maximum parsimony algorithm. Further trees were constructed using 1000 bootstrap iterations with an LG-G substitution model, a gamma distribution with three gamma parameters, and nearest-neighbor-interchange (NNI) as the heuristic method.

### Yeast two-hybrid assay

AH109 yeast cells (Clontech) were used for yeast two-hybrid experiments. Transformation of yeast cells was performed according to the manufacturer’s guidelines using the Frozen-EZ Yeast Transformation Kit (Zymo Research). Genes of interest were amplified from wild-type genomic DNA using Phusion Polymerase (NEB), according to the manufacturer’s guidelines. The indicated restriction sites were introduced using oligonucleotides listed in Table S1A. Vectors and PCR fragments were cut with the respective restriction enzymes, and the gene of interest was ligated into the vector, leading to a fusion protein with a GAL4 activation domain (AD) or GAL4 DNA-binding domain (BD) either at the N-or C-terminus. All constructed plasmids are listed in Table S2B. The detailed protocol for the growth assay can be found in protocols.io (dx.doi.org/10.17504/protocols.io.wcnfave). Successfully transformed cells were selected on a complete supplement mixture (CSM) lacking leucine and tryptophan (-Leu -Trp) dropout medium (MP Biochemicals) at 30°C for 3-4 days. Cells containing bait and prey plasmids were streaked on CSM lacking leucine, tryptophan, and histidine (-Leu -Trp -His) dropout medium (MP Biochemicals) with the addition of 12.5 mM 3-amino-1,2,4-triazole (3-AT, Roth) and incubated for 6 days at 30°C to screen for interactions.

### Expression and purification of recombinant Kai proteins

*Synechocystis* KaiB3, KaiB1 and *Synechococcus* KaiA (plasmids kindly provided by T. Kondo, Nagoya University, Japan) were produced as GST-fusion proteins in *E. coli* BL21(DE3) as described in^25^ (https://www.protocols.io/view/expression-and-purification-of-gst-tagged-kai-prot-48ggztw). Briefly, proteins were purified by affinity chromatography using glutathione-agarose 4 B (Macherey and Nagel), and the N-terminal GST-tag was removed using PreScission Protease (Cytiva) prior to elution of the untagged proteins from the glutathione resin. *Synechocystis* KaiC3 was produced with an N-terminal-Strep-tag (Strep-KaiC3) in *E. coli* Rosetta-gami B (DE3) cells and purified via affinity chromatography using Strep-tactin XT superflow (IBA-Lifesciences)^25^ (https://www.protocols.io/view/heterologous-expression-and-affinity-purification-meac3ae). *The Synechocystis* ORF *sll0485,* encoding KaiA3, was inserted into the vector pET22b to create a C-terminal His6-fusion. KaiA3-His6 was expressed in *E. coli* Tuner (DE3) cells and purified by immobilized metal affinity chromatography (IMAC) using PureProteome™ Nickel Magnetic Beads (Millipore). For a detailed protocol, see at protocols.io (<u>dx.doi.org/10.17504/protocols.io.bu5bny2n</u>). Recombinant proteins were stored at -80°C in buffer containing 20 mM Tris, pH 8.0, 150 mM NaCl, 0.5 mM EDTA, 5 mM MgCl_2_, and 1 mM ATP.

### KaiC3 phosphorylation in *in vitro* assays and liquid chromatography mass spectrometry (LC-MS/MS)

Recombinant Strep-KaiC3 purified from *E. coli* exists mainly in its phosphorylated form (KaiC3-P). Fully dephosphorylated Strep-KaiC3 (KaiC3-NP) was generated by incubating the protein for 18 h at 30°C in assay buffer (20 mM Tris, pH 8.0, 150 mM NaCl, 0.5 mM EDTA, 5 mM MgCl_2_, and 1 mM ATP). The autokinase activity of KaiC3-NP was investigated by incubating 0.2 µg/µl KaiC3 for 16 h at 30°C in 20 µl assay buffer in the presence or absence of 0.1 µg/µl KaiA3-His6, KaiB3, and *Synechococcus* KaiA. Ten-microliter aliquots were taken before and after incubation at 30°C, and the reaction was stopped with SDS sample buffer. Samples were stored at -20°C prior to application to a high resolution LowC SDS gel (10% T, 0.67% C)^84^ using the Hoefer Mighty small II gel electrophoresis system and Tris-Tricine running buffer (cathode buffer: 100 mM Tris, 100 mM Tricine, 0.1 % SDS, pH 8.25; anode buffer: 100 mM Tris, pH 8.9, according to Schägger and von Jagow^85^). Gels were stained with Coomassie Blue R.

For the 48 h assay, pools containing 0.2 µg/µl (3.4 µM) KaiC3-NP, 0.1 µg/µl KaiB3 (7.4 µM) and various concentrations of KaiA3-His6 (corresponding to 0.5 – 8.4 µM) were prepared in assay buffer supplemented with 5 mM ATP, split in 10 µl aliquots for the desired timepoints and stored at -80°C. Samples were thawed on ice for 10 min prior to incubation at 30°C for the different time periods. The reaction was stopped at specific time points by adding SDS sample buffer. Samples were stored at -80°C prior to application to a LowC SDS gel (10% T, 0.67% C)^26^ sing the Biorad Mini PROTEAN gel electrophoresis system and Tris-glycine running buffer (25 mM Tris, 192 mM glycine, 0.1 % SDS, according to Laemmli^86^). The gels were stained with ROTI®Blue quick stain. In Tris-glycine buffer, three KaiC3 bands could be separated, whereas two KaiC3 bands were separated in Tris-Tricine buffer.

For LC-MS/MS-based analysis of KaiC3 phosphorylation sites, Strep-KaiC3 and KaiA3 were co-incubated *in vitro* as described above. Samples were taken directly after mixing, as well as after 2 and 6 h of incubation, and separated by SDS-PAGE. For each sample, protein-containing gel regions of Strep-KaiC size were cut out with a scalpel. For the 6 h time point, a gel region at the potential size of the Strep-KaiC3/A3 complex was also extracted. In-gel protein digestion with trypsin was performed as described by Shevchenko *et al.* ^87^. The generated peptides were extracted and purified using the stage tip protocol^88^. Of the resulting peptide solution, 20% was used for nanoLC-MS/MS analysis. Therefore, peptides were separated in a 37 min reverse-phase linear gradient and directly ionized in an online coupled ESI source upon elution for analysis on a Q Exactive HF mass spectrometer (Thermo Fisher Scientific) operated in data-dependent acquisition mode. The 12 highest abundant multiply charged ions of each full scan were separately fragmented by HCD, and the generated fragment ions were analyzed in consecutive MS/MS scans. Raw data files were processed using MaxQuant software (version 1.5.2.8) and default settings. Phosphorylation of Ser, Thr, and Tyr was defined as a variable modification. Acquired m/z spectra were searched against the proteome databases of *Synechocystis* and *E. coli* (downloaded from cyanobase and Uniprot, respectively). Annotated MS/MS spectra were visualized using the MaxQuant viewer.

### Clear native protein PAGE, Phos-tag SDS-PAGE and immunodetection

Kai proteins (10 µl samples containing 2 µg dephosphorylated Strep-KaiC3, 1 µg KaiA3-His6, 1 µg KaiB3, 1 µg *Synechococcus* KaiA) were incubated for 16 h at 30°C in phosphorylation assay buffer, followed by separation of the native proteins in 4-16% native PAGE at 4°C using a clear native buffer system (Serva) without anionic dye. Thus, only proteins with a pI<7 at physiological pH were separated. Protein bands were visualized with Coomassie staining (ROTIBlue Quick, Carl Roth) or immunodetected with a monoclonal anti-His antibody conjugated to HRP (MA1-21315-HRP, Thermo Fisher, 1:2000 diluted). A detailed protocol can be found in protocols.io (dx.doi.org/10.17504/protocols.io.bu67nzhn).

To analyze *the in vivo* phosphorylation of KaiC3, *Synechocystis* wild type, *ΔkaiA3,* and *ΔkaiC3* cells were cultivated in BG11 or copper-depleted medium for *kaiA3* overexpression. After an initial 12h/12h light/dark cycle, 10 ml of cells were collected every 6 h for analysis. The cells were cooled in liquid nitrogen for 5 s and harvested by centrifugation (3220 × g, 2 min, 4°C). The pellet was frozen in liquid nitrogen and stored at -20°C until further processing. To lyse the cells, the pellets were resuspended to an OD_750_ of 25 in phosphorylation buffer (50 mM NaOH-HEPES pH 7.5, 300 mM NaCl, 0.5 mM Tris-(2-carboxyethyl)-phosphine, 10 mM MgCl_2_). The cells were disrupted twice in a cell mill at 30 Hz for 1 min at 4°C, using glass beads. The crude cell extract was obtained by centrifugation (500 × g, 1 min, 4°C). For mobility shift detection of phosphorylated and dephosphorylated KaiC3, a Zn^2+^-Phos-tag® SDS-PAGE assay (Wako Chemicals) was used. A 9% SDS-PAGE gel containing 25 µM Phos-tag acrylamide was prepared and 12 µL of cell extract was run at 150 V for 3 h at 4°C. Proteins were blotted onto a nitrocellulose membrane (Amersham™ Protran®) via wet blotting. Immunodetection was performed using αKaiC3^27^ and anti-rabbit secondary (Thermo Fisher Scientific Inc., USA) antibodies.

### Screening of KaiC3 and KaiC1 binding partners by immunoprecipitation-coupled liquid chromatography mass spectrometry (LC-MS/MS)

*Synechocystis* WT/FLAG-*kaiC3*, WT/FLAG-*kaiC1*, and WT/FLAG-*sfGFP* (control) strains were cultivated in BG11 medium (100 ml, copper-depleted) and harvested by centrifugation at 6000 × *g* for 10 min at 4°C. According to Wiegard *et al*.^27^, cells were disrupted in a mixer mill, followed by solubilization with n-dodecyl-β-maltoside for 1 h. The supernatant was used for FLAG purification in pull-down assays with Anti-Flag® M2 Magnetic Beads (Sigma-Aldrich), following the manufacturer’s protocol. The resulting elution fractions were loaded onto a NuPAGE™ Bis-Tris Gel and run following the manufacturer’s protocol (Invitrogen). Protein bands were allowed to migrate only a short distance of approximately 10 mm. After staining the gel for 60 min with InstantBlue™ (Expedeon), the protein-containing gel regions were excised. Two independent replicates were produced for each condition (KaiC3, KaiC1, or control pull-down). In-gel protein digestion with trypsin was performed as described above, and the resulting peptide solutions were purified using stage tips. Approximately 20% of the sample was applied for nanoLC-MS/MS analysis as described above on a Q Exactive HF mass spectrometer (Thermo Fisher Scientific) operated in the data-dependent acquisition mode. Raw data of KaiC3 or KaiC1 pull-downs were separately processed using the MaxQuant software (version 1.5.2.8) embedded MaxLFQ algorithm as described by Cox *et al.*^89^. Raw spectra were searched against the proteome databases of *Synechocystis* and *E. coli* (downloaded from cyanobase and Uniprot, respectively) and the bait protein sequences. Significantly enriched proteins were identified by Perseus software (version 1.6.5.0) significance B analysis with a p-value-of 0.01.

### Strains and growth conditions

Wild-type *Synechocystis* (PCC-M, resequenced^28^), the deletion strains Δ*rpaA*^37^, Δ*kaiC3*^25^, Δ*kaiA3*, and Δ*kaiA3B3C3* (Fig. S1), and complementation strain Δ*kaiA3*/*kaiA3* (Fig. S1) were cultured photoautotrophically in BG11 medium^90^ supplemented with 10 mM TES buffer (pH 8) under constant illumination with 75 μmol photons m^-2^ s^-1^ of white light (Philips TLD Super 80/840) at 30°C. Cells were grown either in Erlenmeyer flasks with constant shaking (140 rpm) or on plates (0.75% Bacto-Agar; Difco) supplemented with 0.3% thiosulfate. For photomixotrophic experiments, 0.2% glucose was added to the plates. For chemoheterotrophic growth experiments in complete darkness, *Synechocystis* cells were spotted at different dilutions on BG11 agar plates containing 0.2% glucose and incubated either mixotrophically for three days with continuous illumination or chemoheterotrophically in the dark for 26 days.

### Construction of mutants of the KaiC3 based clock system

To construct the *kaiA3* (*sll0485*) deletion strain, *Synechocystis* wild-type cells were transformed with the plasmid pUC19-Δ*sll0485*. For plasmid construction, PCR products were generated using the oligonucleotides P13-P14 and pUC19 as template, P15-P16 and P19-P20 with genomic *Synechocystis* wild-type DNA as template and P17-25 with pUC4K as template. Homologous recombination led to replacement of the *sll0485* gene with a kanamycin resistance cassette (Fig. S1). For genomic complementation of the Δ*sll0485* strain, cells were transformed with the plasmid pUC19-Δ*sll0485*-compl. Overlapping fragments were generated using the oligonucleotides P15-28 and P24-32 with genomic *Synechocystis* wild-type DNA as template, P13-P26 and pUC19 as template, and P22-P23 and the vector pACYC184 as template. In the resulting complementation strain Δ*kaiA3/kaiA3*, the kanamycin resistance cassette was replaced with *sll0485*, and a chloramphenicol resistance cassette was introduced downstream of the *kaiB3* gene (Fig. S1). For the triple-knockout mutant Δ*kaiA3B3C3,* Δ*kaiC3* cells were used as the background strain for transformation with the pUC19-Δ*kaiA3B3* plasmid. PCR products were generated using the oligonucleotides P13-P26 and pUC19 as template, P17-P27 and pUC4K as template, P15-P16 and P25-P28 with genomic *Synechocystis* wild-type DNA as template. The operon *kaiA3kaiB3* was replaced with a kanamycin resistance cassette (Fig. S1). Complete segregation of the mutant alleles was confirmed using PCR. For the Δ*kaiA3* strain, oligonucleotides P15-P29 were used. Segregation of the complementation strain was confirmed by PCR with P15-P29, P30-P31, and P19-P32. For the triple knockout mutant Δ*kaiA3B3C3,* deletion of the *kaiA3B3* operon was confirmed by PCR using the primer pairs P15-P33 and P19-P30. The *kaiA3B3* chromosomal region of the mutants is shown in Fig. S1.

Ectopic expression of *sll0485* was achieved in wild-type and Δ*sll0485* cells after transformation with plasmid pUR-NFLAG-*sll0485*. The plasmid was constructed via restriction digestion of the vector pUR-N-Flag-xyz, and the PCR product was amplified with the oligonucleotide pair P29-P34 using genomic *Synechocystis* wild-type DNA as a template. Restriction digestion with EcoRI and BamHI was followed by ligation. Successful transformation was confirmed by PCR with P35-P36. The oligonucleotides and plasmids used are listed in Table S1.

## Data availability

The mass spectrometry proteomics data were deposited in the ProteomeXchange Consortium (http://proteomecentral.proteomexchange.org) via the PRIDE partner repository ^91^, with the dataset identifier PXD0042846.

Datasets S1 to S3 were deposited on a server and can be accessed under the following link: https://supplements.biologie.uni-freiburg.de/the_non-standard_kaia3_regulator/

## Information for reviewers

account: pilus, password: freecastle

## Acknowledgments

We thank Research Unit FOR2816 ‘SCyCode’ (The Autotrophy-Heterotrophy Switch in Cyanobacteria: Coherent Decision-Making at Multiple Regulatory Layers) funded by the German Research Foundation for fruitful discussions and financial support. This work was financially supported by grants (WI2014/5-3; 10-1; AX 84/1-3 and MA 4918/4-1) from the German Research Foundation to A.W., I.M.A., and B.M., respectively. We thank Pauline Morys, Annika Klopp, Isabell Bleile, Werner Bigott, and Petra Kolkhof for technical assistance.

## Supplementary Information for

### Other supplementary materials for this manuscript include the following

Datasets S1 to S3 (online only under the following link): https://supplements.biologie.uni-freiburg.de/the_non-standard_kaia3_regulator/ Information for reviewers: account: pilus, password: freecastle

**Fig. S1.**
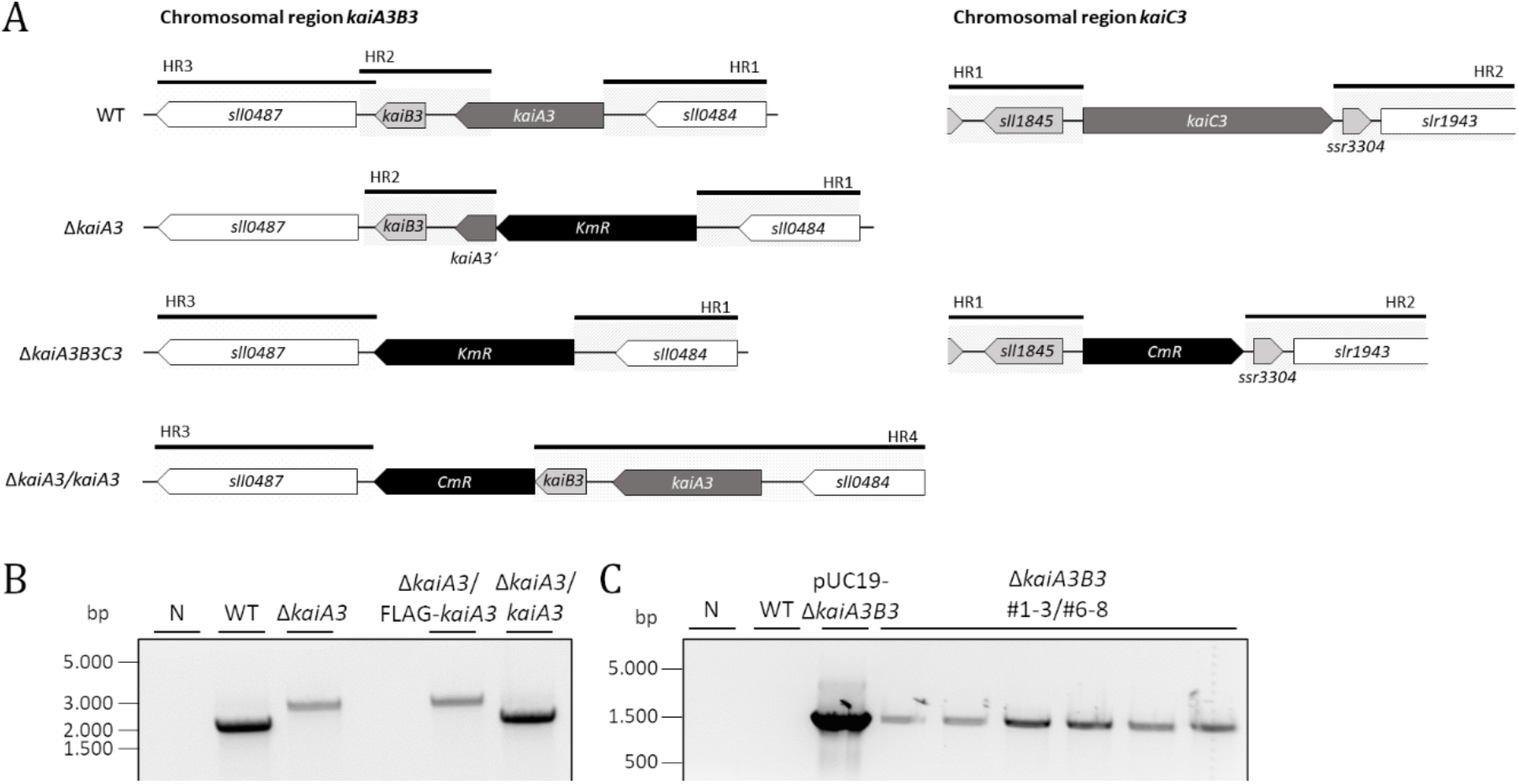
Construction of mutants of the KaiC3 based clock system. (A) Schematic depiction of the *kaiA3B3 and* kaiC3 genomic context. Gene locus of *kaiA3B3* with the up- and downstream located genes. *KaiA3* and *kaiB3* are transcribed as an operon together with *sll0484*, the putative promotor is upstream of *sll0484*. For the inactivation of *kaiA3*, the gene was replaced by a kanamycin resistance cassette (KmR). For the construction of the triple knockout mutant Δ*kaiAB3C3*, the genomic region from *kaiA3* to *kaiB3* was replaced by a kanamycin resistance cassette (KmR) in the Δ*kaiC3* strain. Complementation of the Δ*kaiA3* strain was achieved by the introduction of *kaiA3* within its original genomic context with a chloramphenicol resistance cassette introduced downstream of the *kaiB3* gene. Clones were selected for chloramphenicol resistance and lack of kanamycin resistance. Black bars and grey, dashed boxes represent the regions for homologous recombination into the *Synechocystis* chromosome. (B) Representative result for the verification of the complete segregation of the Δ*kaiA3* deletion strain and Δ*kaiA3*/*kaiA3* complementation strain using colony PCR with the oligonucleotides P15-P29 (Table S1A). A non-template reaction (N), chromosomal WT DNA and Δ*kaiA3*/FLAG-*kaiA3* served as control reactions. Expected construct sizes are 1913 bp for the WT allele and Δ*kaiA3*/*kaiA3*, and 2472 bp for Δ*kaiA3* and Δ*kaiA3*/FLAG-*kaiA3*. (C) Verification of the *kaiA3B3* deletion in the Δ*kaiC3* strain using colony PCR with the oligonucleotides P15-P30. A non-template reaction (N), chromosomal WT DNA and the vector pUC19-Δ*kaiA3B3* served as control reactions. Expected construct size for Δ*kaiA3B3* and pUC19-Δ*kaiA3B3* is 1554 bp. No construct was expected for the WT allele. Complete segregation was verified with the oligonucleotides P19-P30 (not shown).

**Fig. S2.**
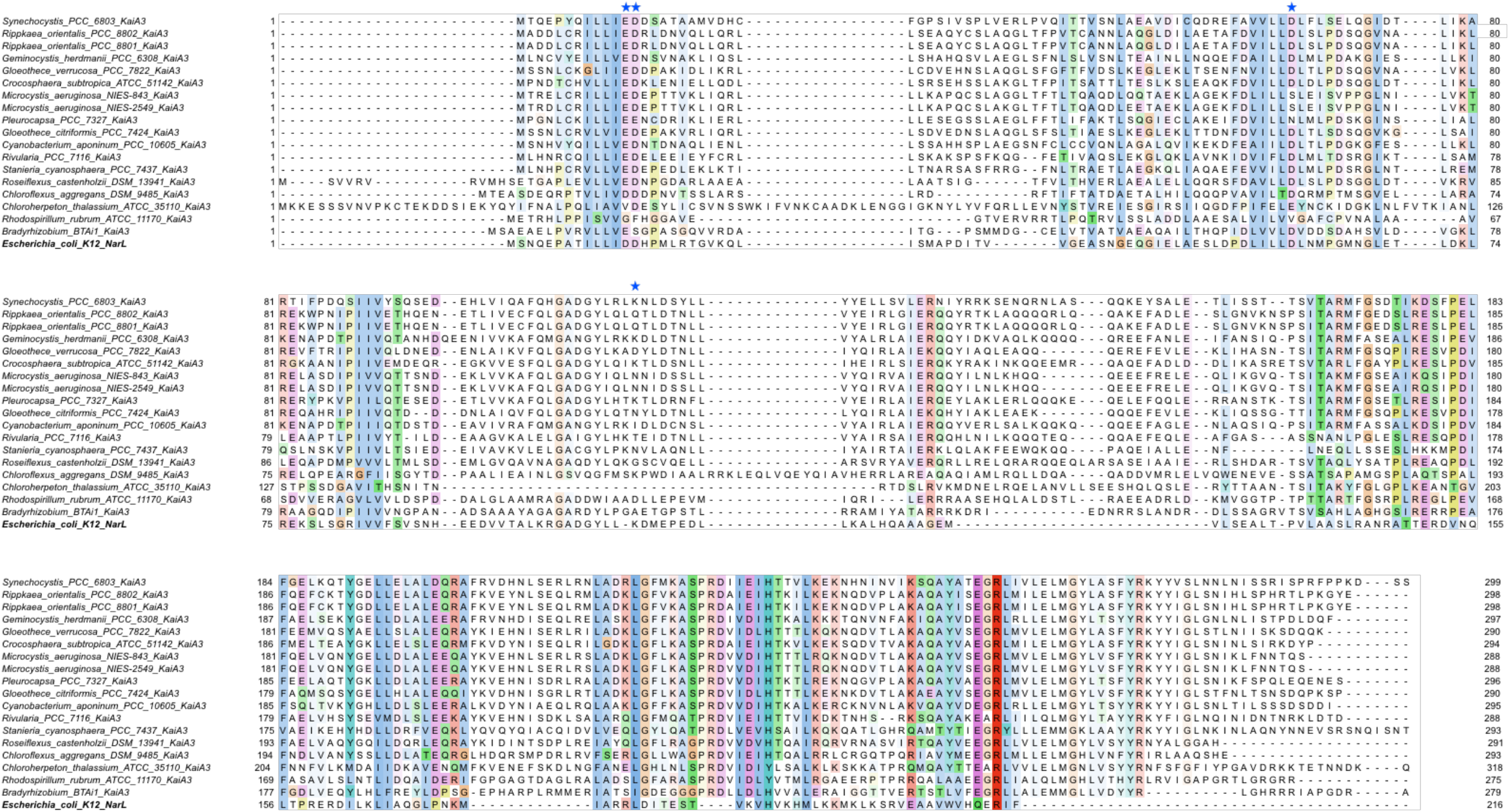
Alignment of the amino acid sequences of Sll0485 (KaiA3) orthologs including NarL from *E. coli*. The sequences were aligned with Mafft (preset, L-INS-i). (A) Sequences are represented in the Clustalx color code with conservation visibility set 20 %^1–3^. As a representative of NarL-type response regulators, the NarL homolog of the *E. coli* strain K12 (UniProtKB - P0AF28) was added. The residues crucial for phosphorylation in response regulators are marked with a blue star^4^.

**Fig. S3.**
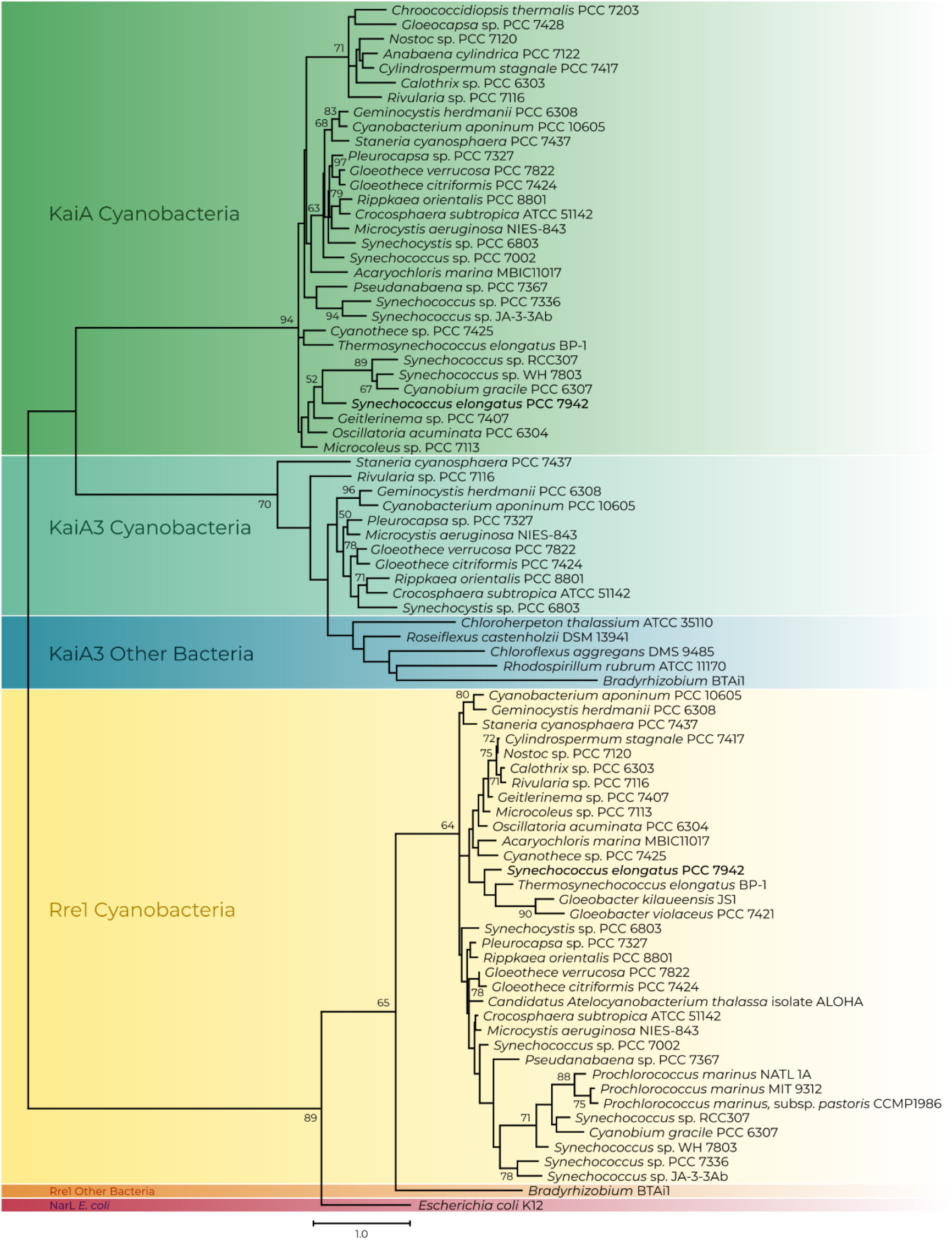
Maximum likelihood-inferred phylogenetic reconstruction of selected orthologs of Sll0485 (KaiA3), Slr1783 (Rre1) and KaiA as well as NarL from *E. coli* (UniProtKB - P0AF28). The sequences were aligned with Mafft (L-INS-i default parameters, Jalview), trimmed to position 168 of the C-terminus of the *Synechococcus elongatus* PCC 7942 KaiA. Aligned sequences were used to infer an unrooted maximum likelihood protein tree. The scale bar indicates 1 substitution per position. Bootstrap values (n=1000) are displayed at branches. Bootstrap values less than 50 are not shown.

**Fig. S4.**
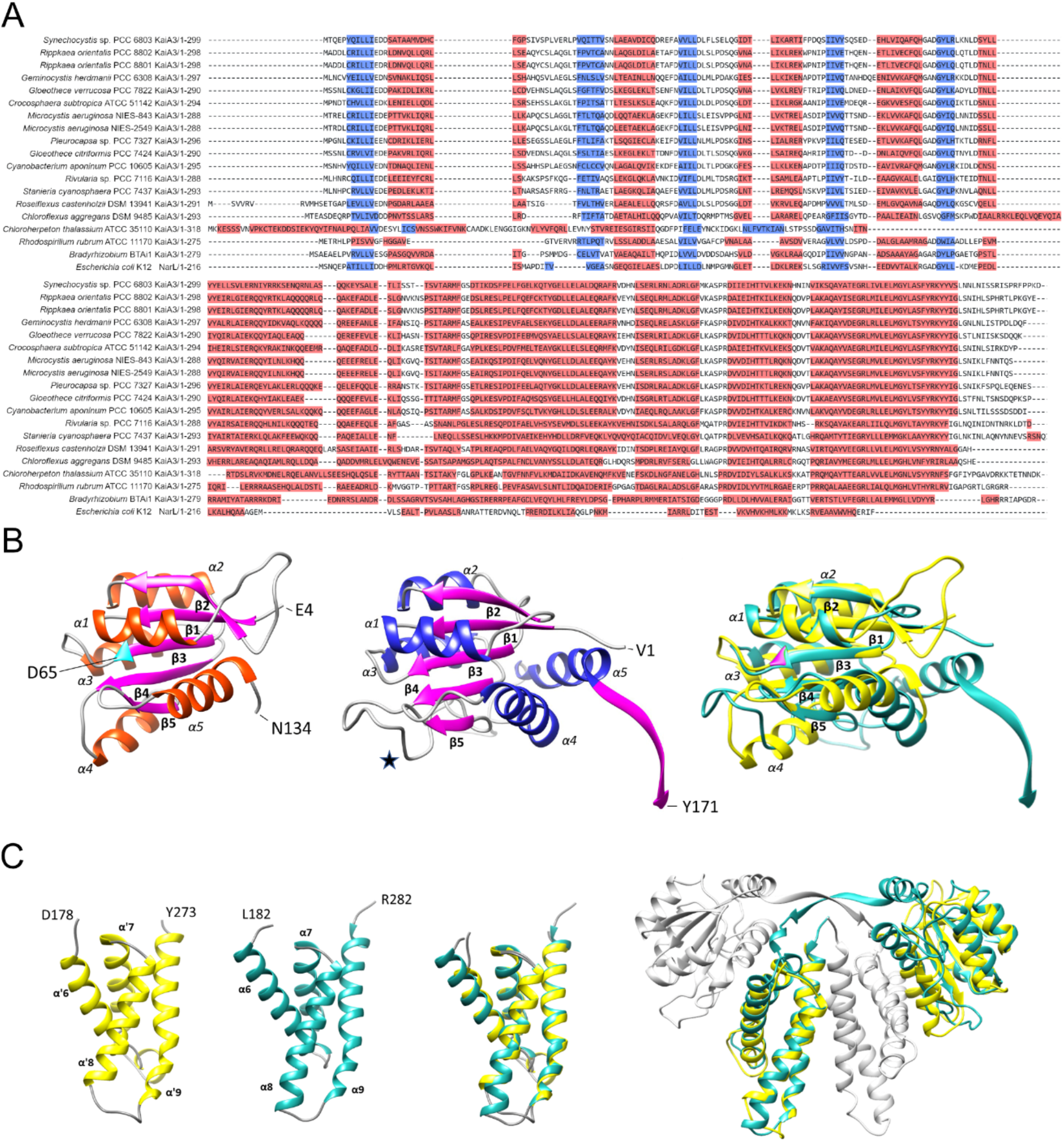
Secondary structure prediction and modelling of KaiA3. (A) Sequence alignment (Fig. S2) was followed by a secondary structure prediction with Ali2D (preset, 30 % identity cutoff to invoke a new PSIPRED run). Predicted α-helices and β-strands are shown in red and blue, respectively^5^; https://toolkit.tuebingen.mpg.de. (B) The structure of the N-terminal domain of KaiA3 is closely similar to the PsR domain of KaiA. Left: The 3-D structure of the KaiA3 N-terminal domain (template *E. coli* NarL, PDB 1A04) was modelled with SWISS-Model (https://swissmodel.expasy.org/). It comprises residues E4-N134 (initial search with residues 1-140) and displays the canonical fold of a response regulator domain: a five-stranded α/β fold with a central five-stranded parallel β-sheet flanked on both faces by five amphipathic α-helices. The predicted phosphorylation site D65 is shown in turquoise. Middle: The PsR domain of *Synechococcus elongatus* PCC 7942 KaiA (template PDB 4G86, residues 1-171 shown) lacks the α4-helix (highlighted by an asterisk) as well as a phosphorylation site. Right: The predicted structure of the KaiA3 N-terminus (shown in yellow) superimposes well on the PsR domain of KaiA (shown in light sea green). The putative phosphorylatable aspartate in the KaiA3 β3-sheet is shown in pink. (C) The structure of the C-terminal domain of KaiA3 displays a KaiA-like motif. Left (yellow): The 3-D structure of the KaiA3 C-terminal domain (template KaiA *Thermosynechococcus elongatus* PDB 1V2Z) was modelled with SWISS-Model (https://swissmodel.expasy.org/). It comprises residues D178 – Y273 (initial search with residues 141-299) and displays a KaiA-like four helix bundle (α’6 - α’9). Left (light sea green): The C-terminal domain of *Synechococcus elongatus* PCC 7942 KaiA (template PDB 4G86, residues 182 - 282). Numbering of the helices according to Ye *et al*.^4^. Middle: Superimposition of the KaiA3 C-terminal domain model structure on the KaiA C-terminus. Right: Superimposition of both KaiA3 domains on the chain B of the KaiA dimer (PDB 4G86). KaiA3 structures are shown in yellow, KaiA structures in light sea green.

**Fig. S5.**
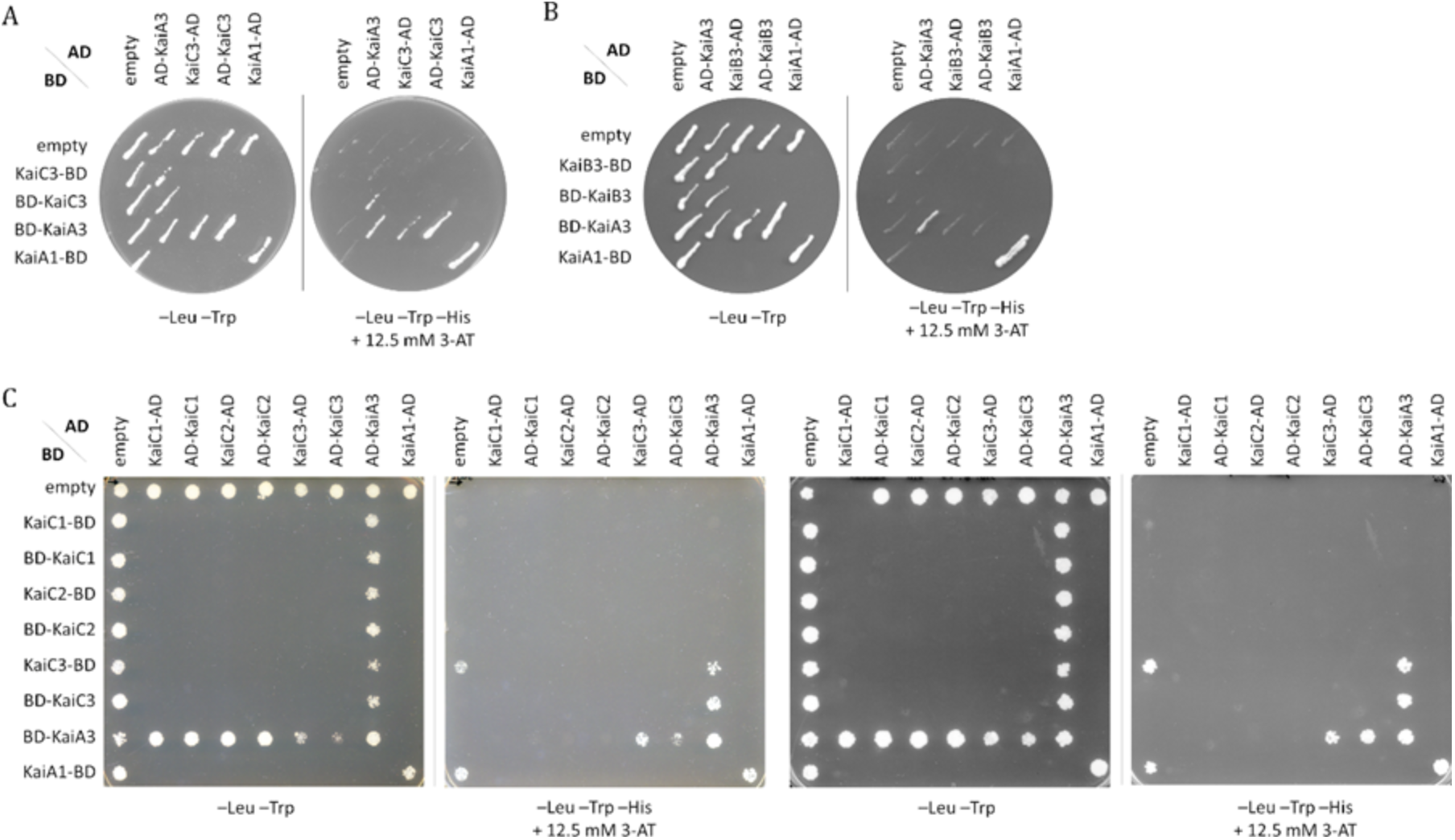
Complete scans of the plates for KaiA3 interaction analysis with KaiB3 and the three KaiC homologs (KaiC1-KaiC3). Yeast two-hybrid reporter strains carrying the respective bait and prey plasmids, were selected by plating on complete supplement medium (CSM) lacking leucine and tryptophan (-Leu -Trp). As a positive control, *Synechocystis* KaiA dimer interaction was used. AD, GAL4 activation domain; BD, GAL4 DNA-binding domain. (A-C) Physical interaction between bait and prey fusion proteins is determined by growth on complete medium lacking leucine, tryptophan and histidine (-Leu -Trp -His) and addition of 12.5 mM 3-amino-1,2,4-triazole 88 (3-AT).

**Fig. S6.**
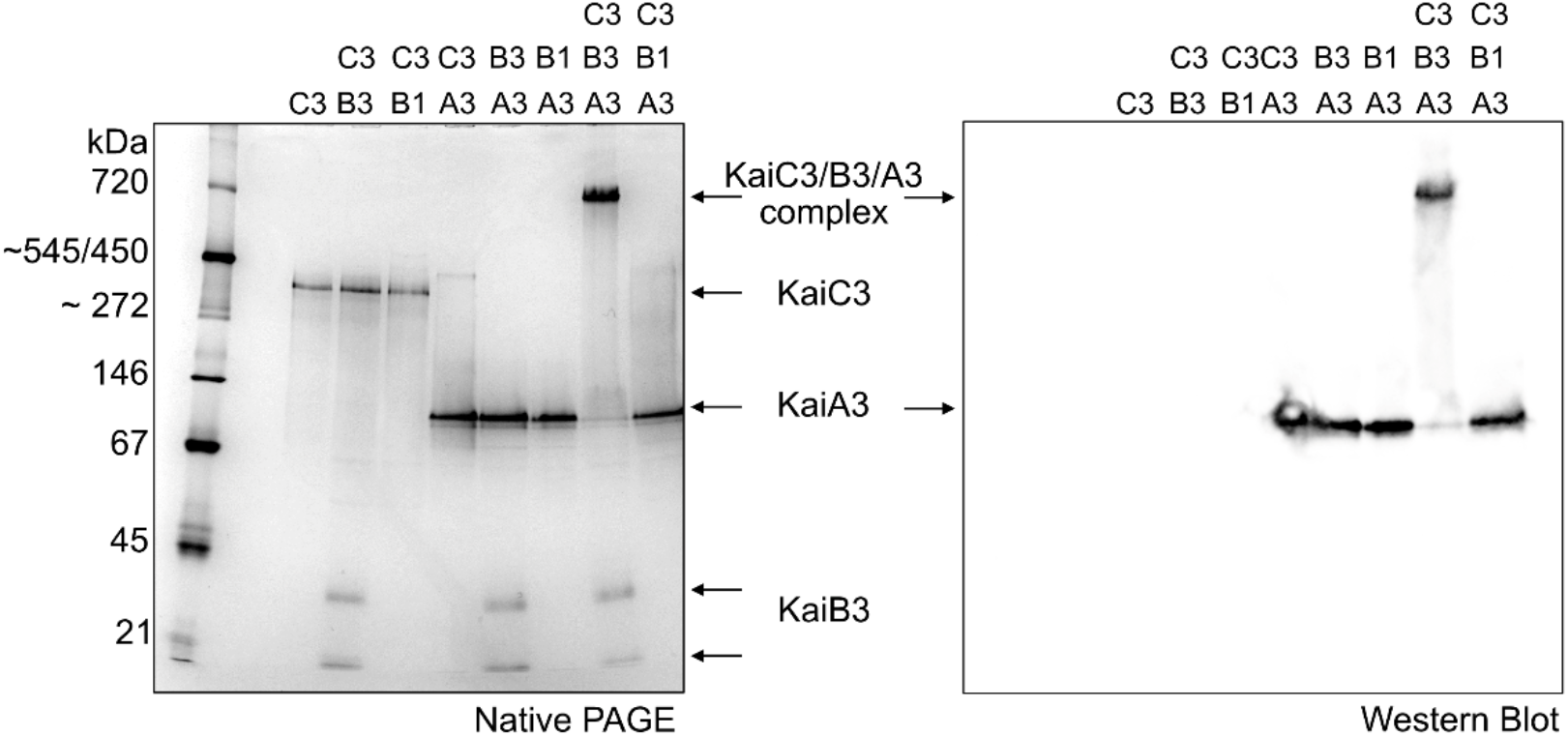
KaiB1 does not form a complex with KaiA3 and KaiC3. Proteins were incubated for 16 h at 30 °C and subsequently subjected to 4-16% clear native PAGE. Gels were either stained with Coomassie Blue (left) or blotted and immunodecorated with a monoclonal anti-His antibody for the detection of recombinant KaiA3-His6 (right). Arrows indicate monomers or protein complexes. The KaiB1 monomer (12 kDa) is not visible in the 4-16% native gradient gel. In contrast to Fig. 2B, a faint band appears in the KaiA3/KaiC3 sample, which is shifted in comparison to the other KaiC3 bands (left). However, KaiA3 is not immunodetected in the shifted band (right).

**Fig. S7.**
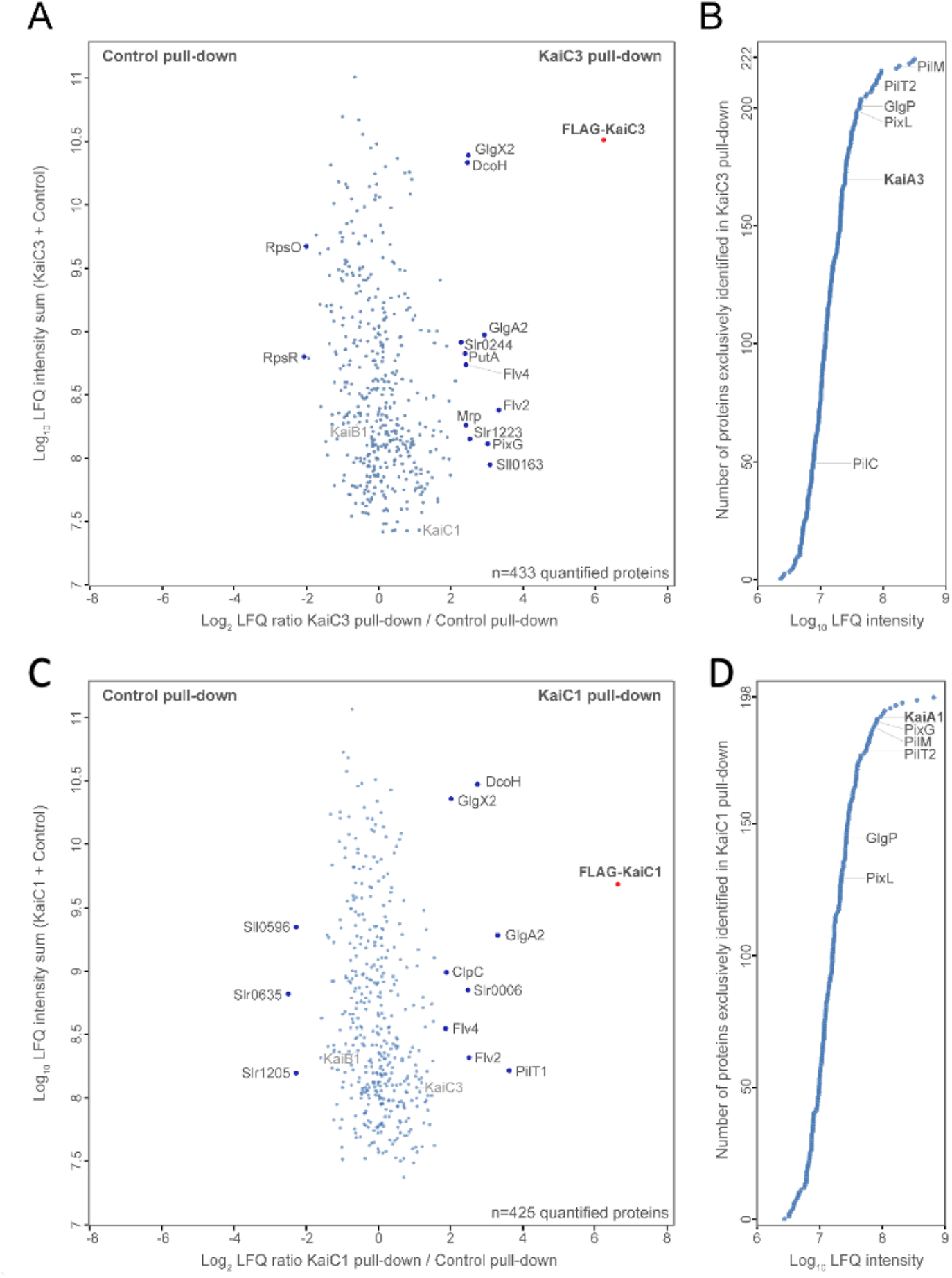
Immunoprecipitation-coupled LC-MS/MS screening of KaiC3 (A, B) and KaiC1 (C, D) binding partners. Solubilized cell lysate of WT/FLAG-*kaiC3* (A), WT/FLAG-*kaiC1* (C) and WT/FLAG-*sfGFP* (control) strains were cultured under continuous light conditions in copper-depleted BG11 medium and used for α-FLAG co-immunoprecipitation in pull-down assays. After FLAG-purification, the elution fractions were analyzed by LC-MS/MS. Label-free quantification using the MaxQuant MaxLFQ algorithm was applied to identify co-enriched proteins. Panels A, C include quantified proteins which were detected in the FLAG-KaiC3 or FLAF-KaiC1 overexpression strain and the control strain. Log_2_ LFQ ratios of FLAG-KaiC / control are plotted against the log_10_ LFQ intensity. Significantly enriched proteins (p-value = 0.01), labeled in dark grey font, are potential interaction partners of KaiC3 or KaiC1. (B, D) Panels include proteins which were exclusively identified in the FLAG-KaiC3 (B) or KaiC1 (D) pull-down, but not in the control. Proteins were sorted by their abundance in the KaiC co-immunoprecipitation eluates and selected proteins were labeled. A full list of identified proteins is shown in Data S2.

**Fig. S8.**
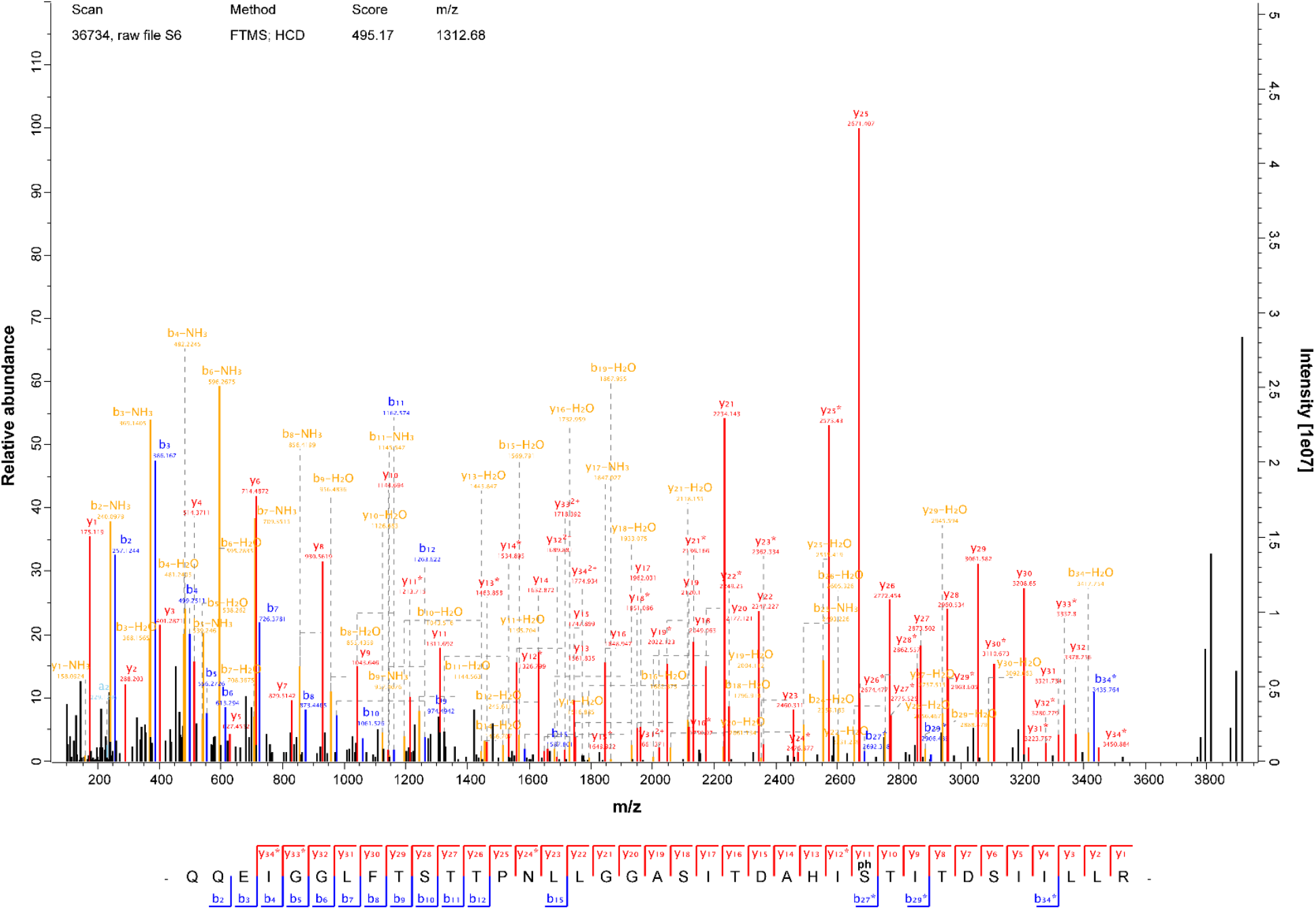
Representative phosphopeptide of KaiC3 detected by mass spectrometry. Samples from KaiC3-KaiA3 *in vitro* co-incubation assays (see materials and method KaiC3 phosphorylation in in vitro assays and liquid chromatography mass spectrometry (LC-MS/MS))were digested with trypsin and analyzed by LC-MS/MS analysis. Comprehensive b- and y-ion series of an abundant, singly phosphorylated 37 amino acid peptide could be detected, localizing the phosphorylation site on Ser423 (position 27 in the peptide). In multiple cases, phosphorylation could be localized on the neighboring Thr424 position instead. Both sites are homologous positions to the KaiC1 auto-phosphorylation sites Ser432 and Thr433 and appeared with increased abundance after prolonged KaiC3-KaiA3 co-incubation duration.

**Fig. S9:**
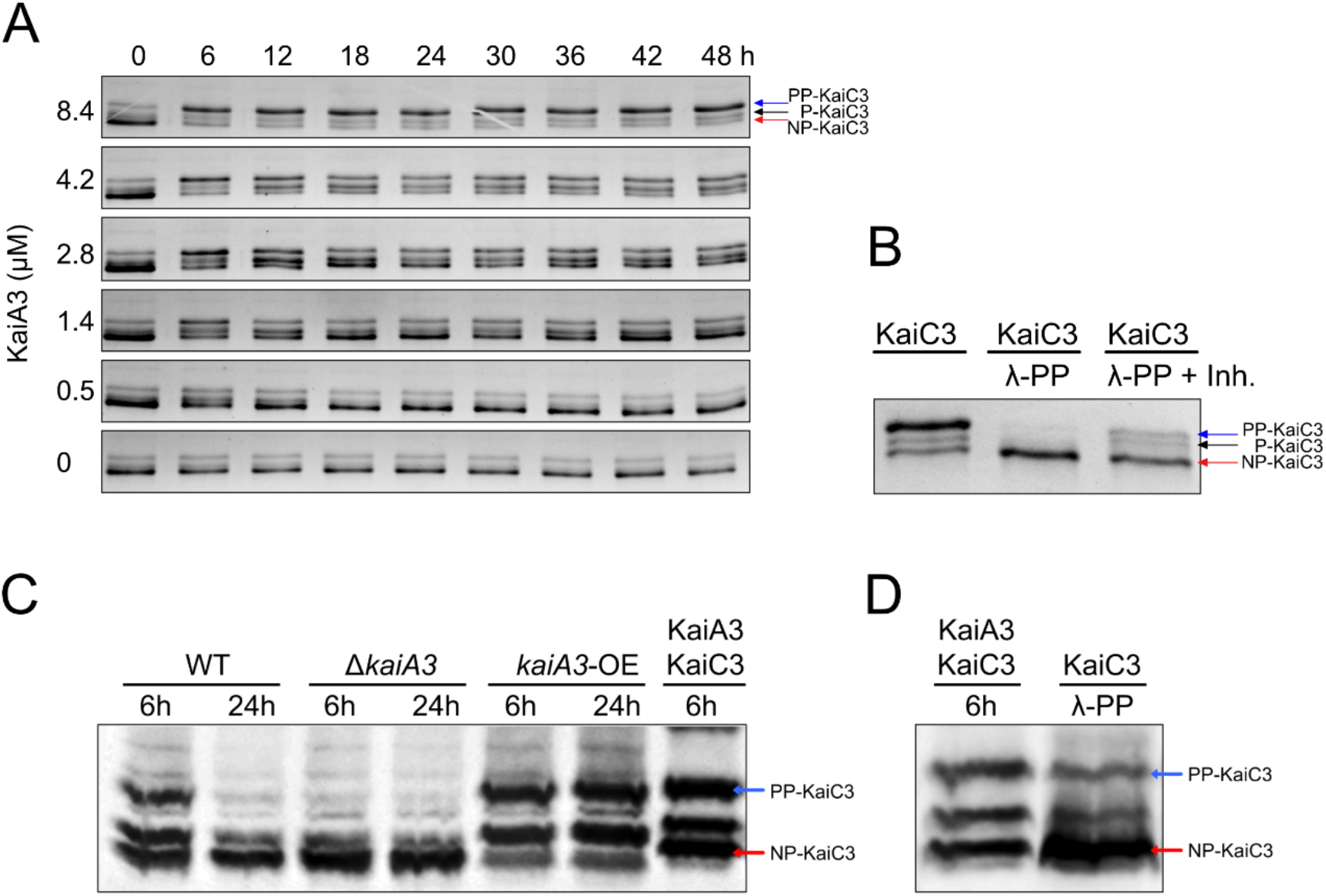
In vitro and in vivo phosphorylation of KaiC3 in dependence of KaiA3. (A) In vitro phosphorylation of KaiC3 in the presence of 7.4 µM KaiB3 and varying concentrations of KaiA3. Representative gel images from 1 assay used for quantification of PP-KaiC3/total KaiC3 displayed in Fig. 3A. (B) KaiC3 was dephosphorylated by incubation with Lambda phosphatase (KaiC3/λ-PP) for 18h at 30°C and separated via high-resolution LowC SDS-PAGE as in A. As control, Lambda-phosphatase activity was blocked by addition of PhosSTOP (Roche) and 10 mM vanadate (KaiC3/λ-PP +Inh.). (C) Comparison of in vivo and in vitro phosphorylation of KaiC3. Whole cell extracts of *Synechocystis* wild-type (WT), *kaiA3* mutant (*ΔkaiA3*), and the overexpression (*kaiA3*-OE) strain, grown in a 12-h light/dark cycle, were subjected to Phos-tag SDS-PAGE followed by western blot analysis with a KaiC3-specific antibody. According to the data shown in Fig. 3C, KaiC3 is in a highly phosphorylated state at 6 h and mostly dephosphorylated at 24 h. For comparison, purified and in vitro phosphorylated KaiC3 were applied to the same gel. (D) KaiC3 was dephosphorylated (KaiC3/PP) using Lambda phosphatase (NEB), applied to a Phos-tag SDS-PAGE and Western blot analysis alongside with in vitro phosphorylated KaiC3, confirming that the fast migrating band of KaiC3 represents the dephosphorylated form of KaiC3. KaiC3 was phosphorylated in vitro in a mixture with KaiA3 (4.2 µM) for 6 h at 30°C (KaiA3-KaiC3/6h).

**Fig. S10.**
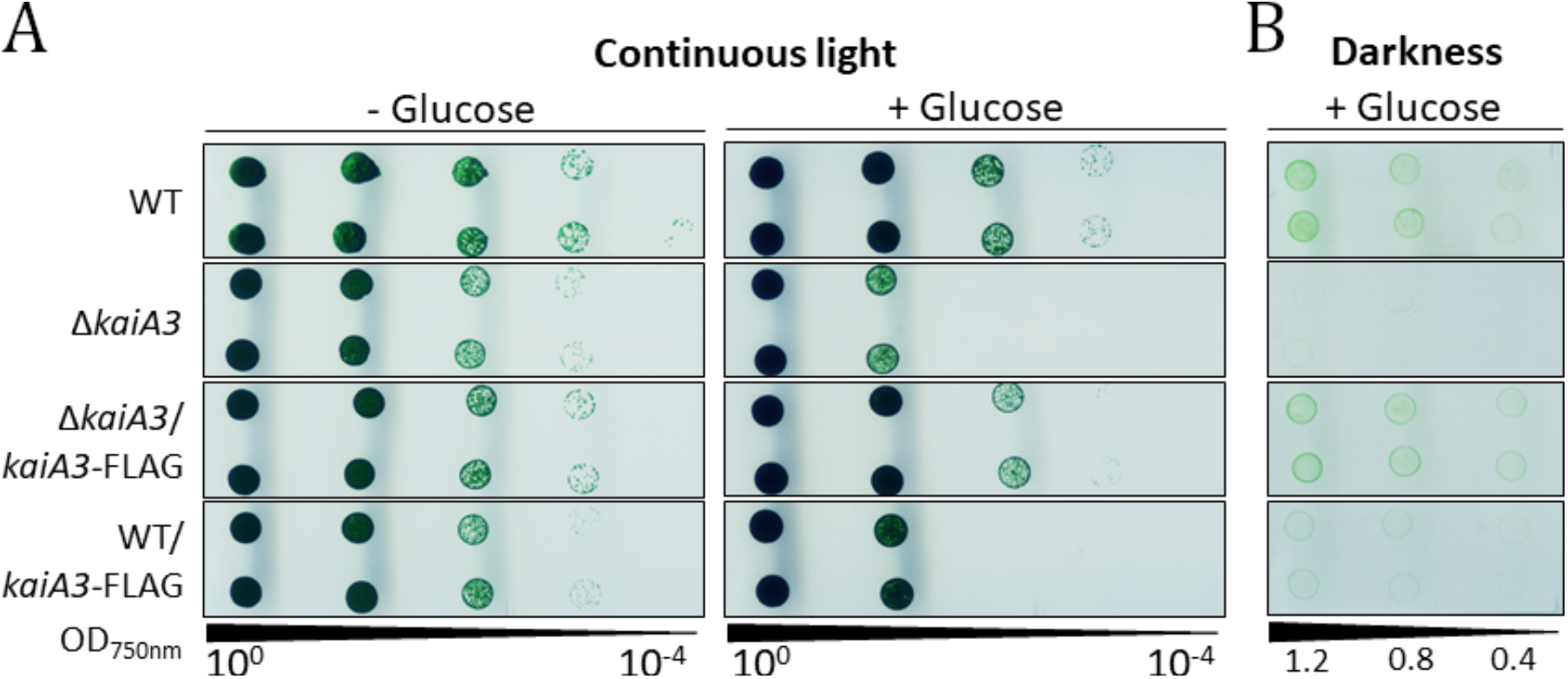
Overaccumulation of *kaiA3* results in growth defects during mixotrophic and chemoheterotrophic growth. Proliferation of the WT, the *kaiA3* deletion mutant, and the strains Δ*kaiA3*/*kaiA3*-FLAG *and* WT/*kaiA3*-FLAG, expressing *kaiA3* ectopically from a self-replicating plasmid, was tested under phototrophic (continuous light, - glucose), photomixotrophic (continuous light, + glucose) and heterotrophic (darkness, + glucose) conditions. Strains were grown in liquid culture under constant light, different dilutions were spotted on agar plates and incubated in the indicated conditions with 75 µmol photons m^-2^ s^-1^ white light (A) or in darkness (B). A representative result of three independent experiments is shown. (A) Cultures were diluted to OD_750nm_ value 0.4 and dilution series were spotted on agar plates with or without the addition of 0.2% glucose. Plates were analyzed after 6 days of continuous light. (B) Cultures were diluted to OD_750nm_ values of 1.2, 0.8 and 0.4 and spotted on agar plates supplemented with 0.2% glucose. Plates were analyzed after 26 days of darkness. For expression of the *kaiA3*-FLAG gene from the P*_petJ_* promoter in the overexpressor strains, all experiments were performed in medium lacking copper.

**Table S1.**
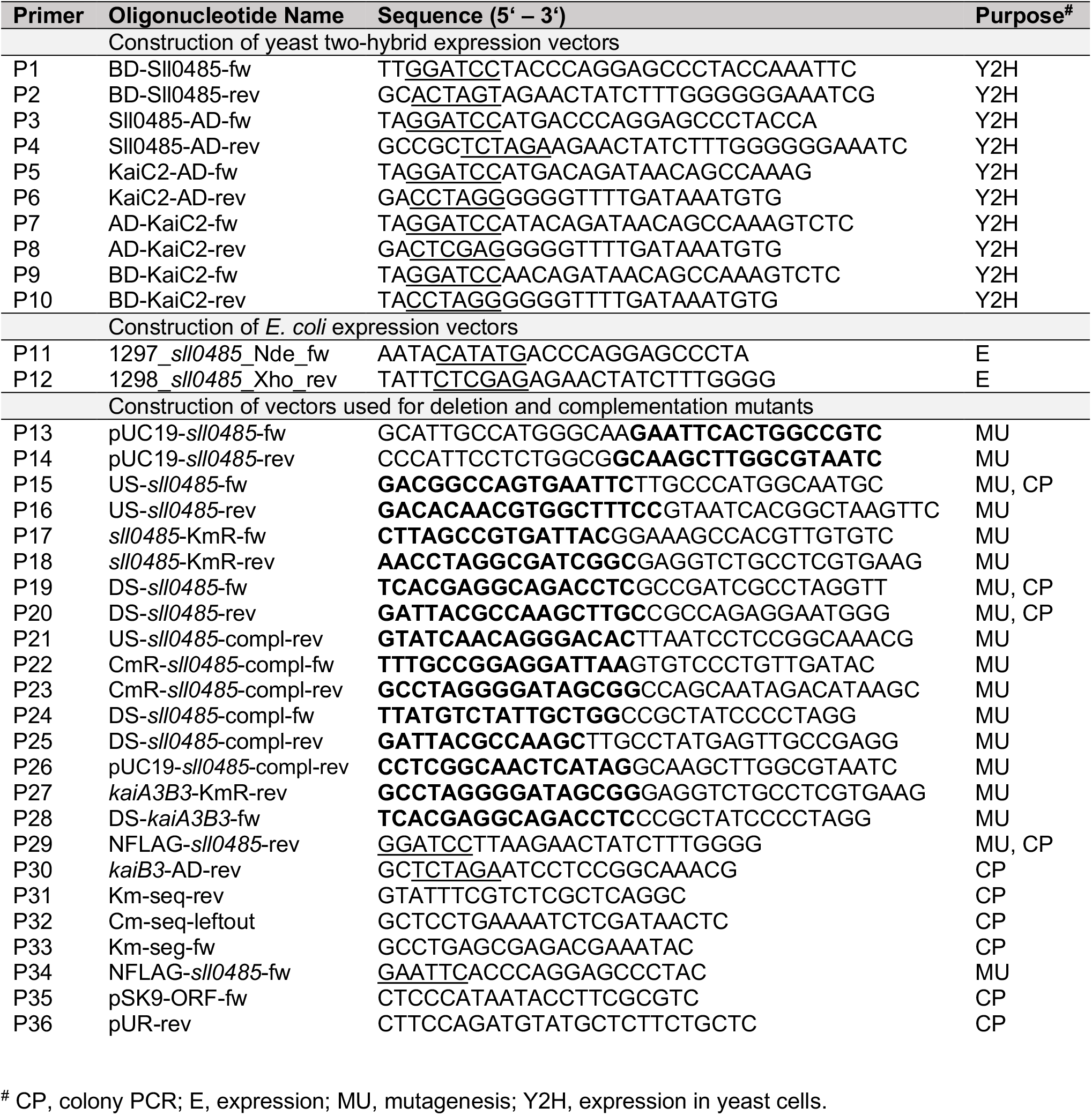
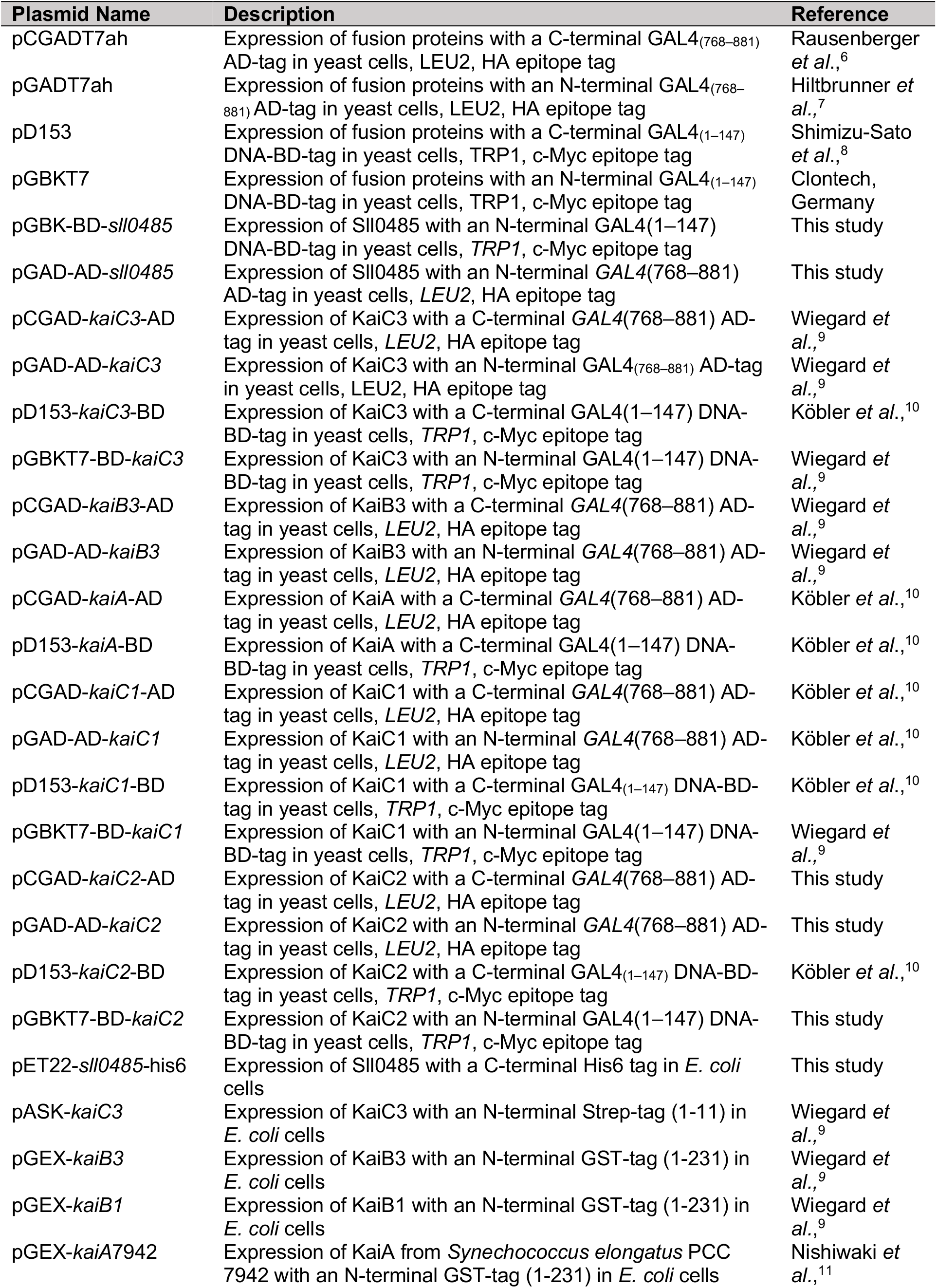
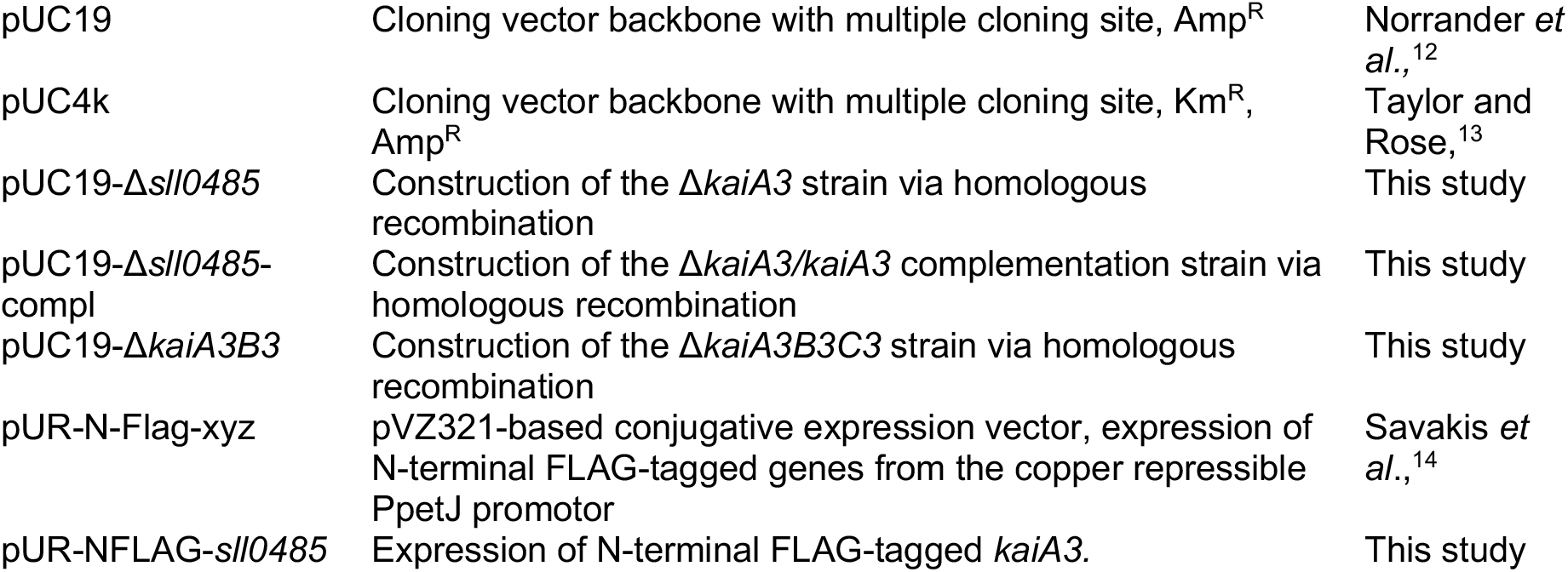
A. Oligonucleotides used in this study. Restriction sites are underlined. Overlaps used for aqua cloning are marked in bold.

**Dataset S1 (separate Excel file).** Putative orthologs of KaiA3 in cyanobacteria and prokaryotes. Header names are described in the following and the exact name is mentioned in parenthesis. Information is provided about the organism (name), the corresponding genus (genus), the taxonomy (taxonomy), and the taxonomic identifier (taxid). Furthermore, the annotated protein name on NCBI (protein), the protein identifier on NCBI (protein_id), the genome identifier where the protein originated from (genome_id), the date when it was last modified on NCBI (date), BLAST statistics (e_value, bitscore, identity), the length of the protein (length) as well as the sequence (seq) were recorded. In addition, the protein id of backward best hit from *Synechocystis* (synechocystis_prot_id) as well as the genome identifier for the genome assembly (synechocystis_id) was stored.

**Dataset S2 (separate Excel file).** Dataset from immunoprecipitation-coupled LC-MS/MS analyses of KaiC3 and KaiC1 interactome analyses. Identified and quantified proteins from label-free analysis of α-FLAG-KaiC3 or -KaiC1 and control co-immunoprecipitation are listed.

**Dataset S3 (separate Excel file).** Dataset of KaiA3B3C3 *in vitro* co-incubation assays on KaiC3 phosphorylation. Localized KaiC3 hphosphorylation sites and phosphorylation occupancies of Ser423/Thr424 are listed.

